# A continuous epistasis model for predicting growth rate given combinatorial variation in gene expression and environment

**DOI:** 10.1101/2022.08.19.504444

**Authors:** Ryan M. Otto, Agata Turska-Nowak, Philip M. Brown, Kimberly A. Reynolds

## Abstract

The ability to predict changes in cellular growth rate given variation in gene expression is critical to engineer biosynthetic pathways and understand evolutionary constraints on mRNA abundance. However, the relationship between gene expression and growth rate is nonlinear and shaped by environmental and genetic context. To address these challenges, we examined the relationship between enzyme expression and *E. coli* growth rate for 36 metabolic gene pairs by measuring growth following 4,404 pairwise, titrated changes in enzyme abundance under varied environmental contexts. Using 20% of these data, we trained an interpretable epistatic model to predict growth rate following simultaneous expression and environmental perturbations. The model’s predictions were robust to significant genetic and environmental epistasis and at times approached the accuracy of biological replicates. Based on these results, we propose a strategy using sparsely sampled, low-order measurements to quantify genetic interaction landscapes on the pathway-wide or genomic scale in a single assay.

## Introduction

Variation in gene expression tunes organism-wide traits like biomass yield, cell division, and differentiation, but predicting these broad phenotypes based on molecular data remains challenging. In the context of bacterial infection, expression changes can lead to immune evasion^1^, trigger a switch between commensal and pathogenic behavior^2^, or confer antibiotic tolerance or resistance^3, 4^. Similarly, dysregulated transcription in human cells can initiate tumorigenesis and metastasis^5, 6^ and transcriptional diversity within tumors confers differential susceptibility to chemotherapeutics and radiation^7, 8^. If transcriptional molecular markers can be leveraged to predict these phenotypes, we could drastically improve patient prognosis. More generally, the relationship between gene expression and cellular growth rate represents a well-defined instantiation of the broader problem of mapping genotype to phenotype.

However, two complications stand in the way of making such predictions. First, expression changes occur over a continuous range and are often non-linearly related to growth rate^9–11^. Thus, a simple on/off mechanism is rarely sufficient to describe the continuum of expression levels observed in a population. While “endpoint” studies such as gene knockouts or near-complete knockdowns have classically proved useful for annotating gene function^12^, discovering new regulatory networks^13^, and revealing the organization of the cell^14–17^, they do not offer sufficient information to predict the effects of intermediate changes in gene expression. Sampling a range of perturbation strengths is required to quantify this continuous, nonlinear response. Second, expression-growth rate relationships are context-dependent^9, 10, 18^. An organism’s genetic background and environment may alter the effect of gene expression changes, a phenomenon called epistasis^19, 20^. The amount of data required to densely sample these context-dependent relationships scales exponentially with the complexity of the system. As a consequence, these experiments quickly become intractable. A new approach is required to approximate these landscapes using a reasonable amount of input data.

Interestingly, prior work on a related problem – predicting the effects of antibiotic and chemotherapeutic combination therapies on cell growth – has considered continuous and/or context-dependent models^21–28^. Of particular relevance is the dose-response model introduced in Zimmer et al., 2016^28^. The authors examined cases where the presence of one drug altered the effect of a second drug. This interaction resulted in a change in the “effective dose” of the second drug and was summarized with two coupling constants – the effect of drug “A” on “B”, and the effect of drug “B” on “A”. The authors found that a model constructed using only these pairwise coupling terms could predict the effects of higher-order combinations of drugs. This model was unit-agnostic, meaning the framework could be trained on any quantitative input, not just drug treatment. However, to our knowledge, it has never been extended to make predictions of growth rate given variation in gene expression or nutrient abundance. Following the rationale that a reduction in enzyme expression is functionally analogous to inhibiting an enzyme with a drug, as well as studies identifying transcription as the main determinant of protein abundance in bacteria^29^, we hypothesized that a similar coupling-based approach could capture context-dependence between genes and predict the impact of multiple gene expression changes on bacterial growth rate.

In this work, we considered continuous expression-growth rate relationships for *E. coli* metabolic enzymes drawn from diverse metabolic pathway architectures. We measured the growth rate effects of 4,404 pairwise combinations of titrated gene knockdowns. Using 20% of these data, we fit a continuous, coupling-sensitive model that was able to predict growth rates for the remaining 80% of pairwise perturbations. We then tested the model’s ability to predict the effect of third-order gene knockdowns without being trained on high-order data and found that sparse pairwise interaction measurements were sufficient to recapitulate the phenotypes observed in this high-order study. Finally, we extended this model to predict the combined effect of gene expression and varied media additives. Growth rate predictions for third-and fourth-order genetic and environmental perturbations remained robust. Throughout these studies, this interpretable approach provided biological insight by identifying asymmetrical genetic and environmental interactions as well as hinting at the existence of high-order interactions in regimes where experimental data diverged from the model’s predictions. Taken together, our experimental and computational methods provide a strategy to incorporate sparsely sampled epistasis measurements into a predictive model. This approach can be used to predict bacterial growth rate following numerous intermediate gene expression changes in a variety of environments and is scalable to quantify expression-growth rate coupling across entire metabolic pathways or potentially even genomes.

## Results

### Growth rate titrates with gene expression following CRISPR interference

Bacterial gene expression is a key determinant of cellular growth rate, and gene expression levels are constrained at both absolute and relative levels. Within these constraints, however, tuning of gene expression can modulate, but not eliminate, bacterial growth (Fig 1A). Classic genetic interaction measurements consider pairwise interactions between genetic knockouts or strong knockdowns and do not permit prediction of growth rates following intermediate changes in expression or gene product activity (Fig 1B). We sought to capture this intermediate variation by generating a continuous expression-growth rate model for a group of metabolic genes in *E. coli* (Fig 1C). However, the amount of data required to directly map these epistatic landscapes scales exponentially as you consider additional expression levels or new genes. Thus, our modeling strategy needs to predict these continuous epistatic landscapes using only sparsely sampled pairwise measurements.

**Figure 1.**
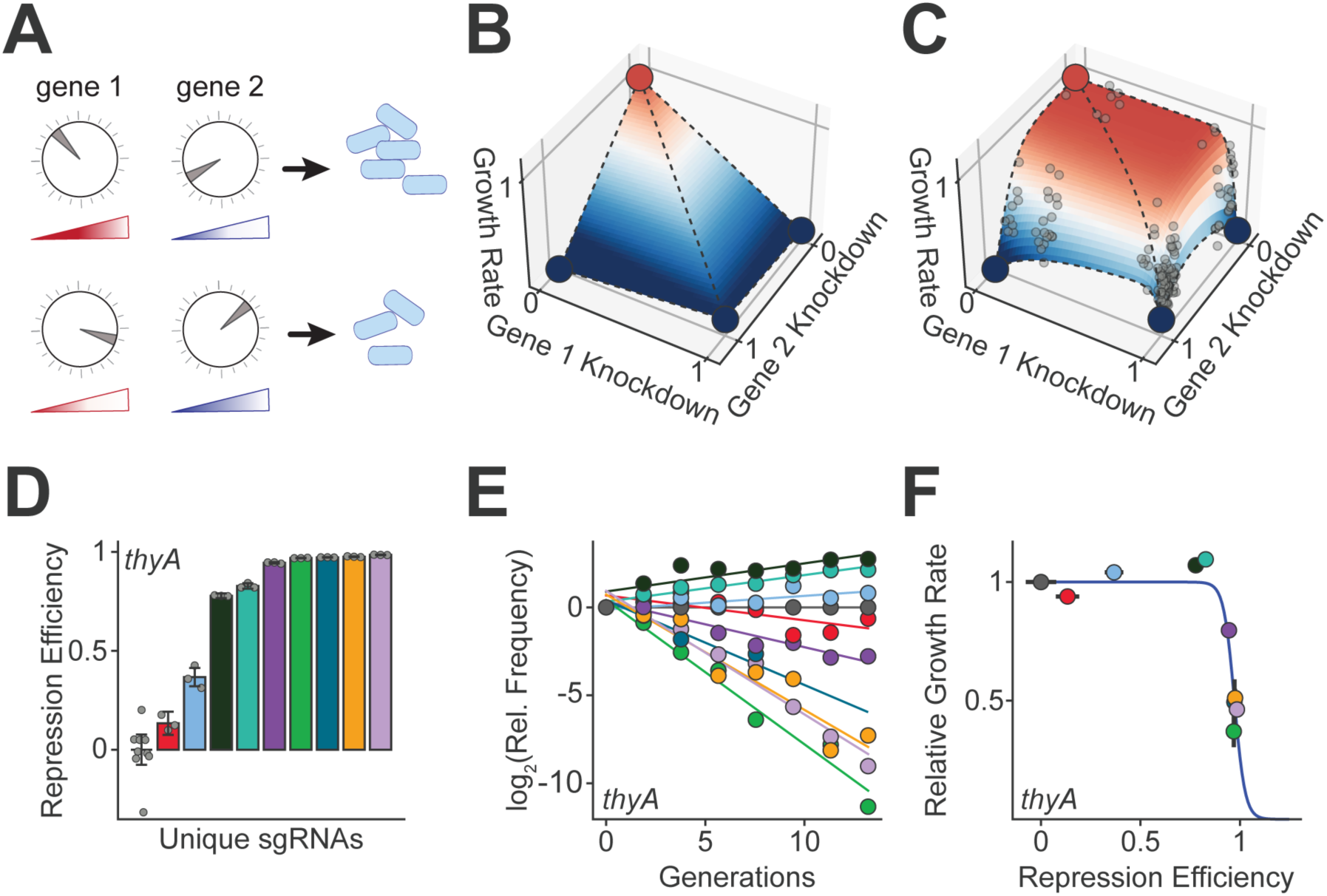
Mismatch CRISPRi enables characterization of gene expression-growth rate relationships. (A) Tuning the expression of *E. coli* genes (dials) and thus relative protein abundances (gradients) alters cellular growth rate. (B) A linear interpolation from experimental data exploring the growth rate effects of complete knockdown of one or two genes, akin to a classic genetic interaction measurement. Growth rate of wildtype cells (x = y = 0, red) linearly decreases with either or both knockdowns of genes 1 and 2 (dotted lines). (C) A computationally modeled, continuous, pairwise expression-growth rate model, fit from intermediate experimental data (gray dots). Growth rate no longer scales linearly with knockdown and is far more robust to expression perturbation than a linear model would suggest. (D) The concentration of mRNA in log-phase *E. coli* for each titrating sgRNA was quantified by RT-qPCR. A repression efficiency of 1 corresponds to complete knockdown. Individual colored bars represent different titrating sgRNAs targeting *thyA,* with error bars describing the standard error of the mean (SEM) across technical replicates. Individual measurements are shown as gray dots. (E) Growth rate was quantified by next-generation sequencing. We measured the log_2_ relative (Rel.) frequency of each sgRNA relative to a non-targeting control (gray dots along y = 0) over eight time points spanning 14 hours. Color coding of sgRNAs is identical to panel D. The slope of the line of best fit represents the growth rate of each knockdown relative to the non-targeting control. (F) The measurements of (rescaled) relative growth rate and repression efficiency show a sigmoidal relationship (blue line), with significant growth rate deficits emerging at severe repression levels. Color coding of sgRNAs is identical to panel D; individual points reflect average relative growth rate across n ≥ 4 replicates, and error bars represent SEM of growth rate measurements (vertical) and RT-qPCR measurements (horizontal).

Training and testing an expression-growth rate model requires high-throughput growth rate measurements across cells harboring distinct, intermediate variations in gene expression. To generate these data, we used mismatch CRISPR interference (CRISPRi). CRISPRi uses a catalytically dead Cas9 endonuclease (dCas9) and a single guide RNA (sgRNA) to target a specific DNA locus, sterically repressing transcription at the target site^30^. In standard CRISPRi, the target of the sgRNA is specified by a 20-nucleotide homology region designed to be fully complementary to the target gene. In contrast, mismatch CRISPRi uses sgRNAs containing nucleotide mismatches in the target homology region^11, 31–33^. These mismatches disrupt sgRNA-DNA interactions, decreasing gene repression strength. By designing a group of sgRNAs targeting the same DNA locus with different numbers of mismatches, we can tune a single gene’s transcription to numerous intermediate levels (Fig 1D). Growth rate defects caused by these knockdowns can be quantified at high throughput using CRISPRi-seq^18, 31, 34–36^. In our implementation of this approach, we transform a library of sgRNAs into *E. coli* containing a chromosomally encoded dCas9 under control of a tet-inducible promoter. To control population size and environmental conditions, we grow the transformed cells in a turbidostat^37^. We then induce each cell’s CRISPRi machinery and use next-generation sequencing to monitor changes in the frequency of individual sgRNAs over time. A linear regression fit to the logarithm of each sgRNA’s frequency over time allows us to estimate the growth rate effects of thousands of sgRNAs in a single experiment (Fig 1E). By plotting growth rate as a function of sgRNA repression strength, we can define continuous expression-growth rate functions for genes of interest (Fig 1F).

In this study, we selected nine genes to target with mismatch CRISPRi, with an eye to sampling distinct metabolic pathways and architectures (Fig 2A-D). Included were genes encoding enzymes catalyzing consecutive metabolic steps (*dapA*/*dapB*, *purN*/*purL*), parallel reactions (*gdhA*/*gltB*), and a three-enzyme loop (*folA*/*glyA*/*thyA*). All genes except *purN*, *gdhA*, and *gltB* were previously shown to be essential in our experimental conditions^38^ (OD_600_ < 0.01 after 24 hours in minimal media). Double knockout of *gdhA*/*gltB* was shown to be synthetically lethal^39^. We designed 9-12 distinct sgRNAs targeting each gene as well as a non-targeting sgRNA control to generate a 96 sgRNA library (Table S1). We quantified the effect of each CRISPRi perturbation on the mRNA concentration of its target gene using reverse transcription-quantitative PCR (RT-qPCR). As expected, our library of sgRNAs created a titrated range of knockdowns for every gene in our data set.

**Figure 2.**
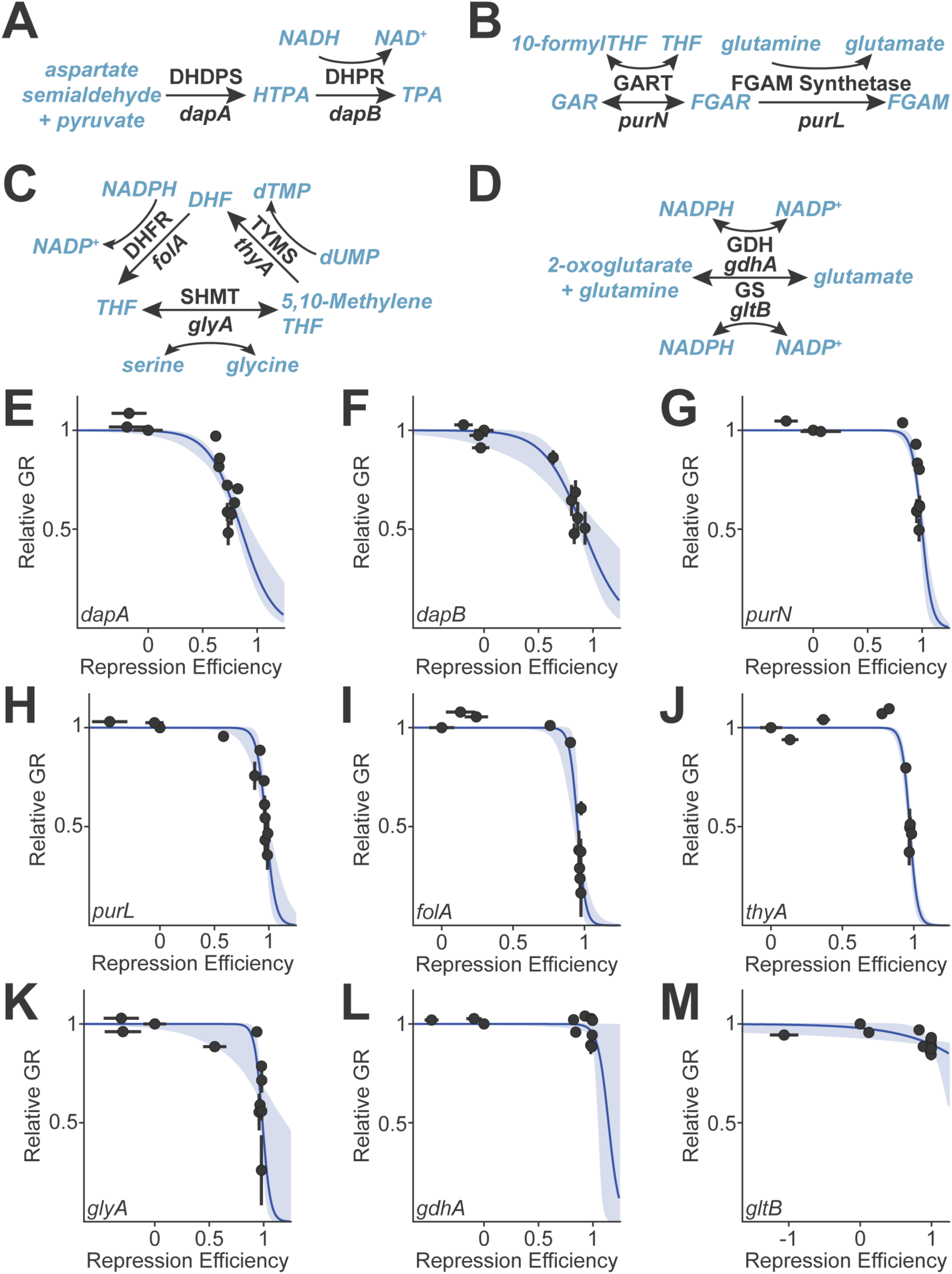
Mismatch CRISPRi for a diverse set of metabolic enzymes. sgRNAs were designed to target 9 *E. coli* genes from (A) lysine, (B) purine, (C) folate, and (D) glutamate metabolism. Gene names are listed in black italic text underneath the corresponding metabolic enzyme (bold black text). Metabolite names are in blue. (E-M) Repression efficiency of each CRISPRi sgRNA, as measured by RT-qPCR, correlated with relative growth rate (GR), as measured by next-generation sequencing. Error bars represent SEM of growth rate measurements between n ≥ 4 replicates (vertical) and RT-qPCR measurements across n ≥ 3 technical replicates (horizontal). Blue lines represent the two-parameter logistic fit to each data set. Blue shaded area represents a pointwise 95% confidence interval of the logistic fit estimated by bootstrapping. The genes *gdhA* and *gltB* are nonessential in our experimental conditions and show no significant growth rate defect following repression.

### Pairwise CRISPRi yields an expression-dependent epistatic landscape

To investigate expression-growth rate coupling between genes, we used Golden Gate cloning to assemble a library of all pairwise combinations of sgRNAs^40^. We constructed this library in three distinct barcoded plasmid vectors to generate internal replicates. Sequencing of our library prior to selection showed that it was complete and well-distributed across the designed sgRNAs (Fig S1A). We quantified the growth rate effect of each CRISPRi knockdown (both singles and pairwise) on exponentially growing *E. coli* cells in M9 minimal media with glucose using continuous culture and next-generation sequencing. Growth rate was well-correlated between internal replicates (Fig S1B). In agreement with earlier work^30^, changing the sgRNA order within pairwise constructs (sgRNA1-sgRNA2 vs. sgRNA2-sgRNA1) did not significantly alter growth rate (Fig S1C). Thus, after pooling our three internal replicates for each of two sgRNA orders, we had six independent measurements for each pairwise CRISPRi treatment. As in prior work^33^, we used these replicates to identify and filter out “escapers”: replicates whose growth rate was significantly faster than all other replicates of the same CRISPRi treatment, likely due to evasion of the CRISPRi machinery (Fig S1D). Escaper correction removed 182 measurements (0.7% of the data, Fig S1E-F). Finally, we averaged the remaining replicates and empirically rescaled our growth rate data, setting the minimum growth rate observed to 0 and the non-targeting CRISPRi control to 1. In total, we quantified the growth rate effect of 4,404 unique CRISPRi perturbations. We found that growth rates from this study correlated well with measurements from our previous work investigating single-gene knockdowns^33^ (Fig S1G).

We quantified single gene expression-growth rate relationships by relating target gene repression strength to growth rate for sgRNA constructs consisting of one targeting and one non-targeting control sgRNA (Fig 2E-M). All genes showed monotonic expression-growth rate relationships except *gdhA* and *gltB*, which were known to be nonessential in our growth condition. Notably, our approach to sgRNA library design reliably sampled the steep, intermediate growth regime of each gene, allowing us to precisely identify the knockdown level where each gene’s transcription became growth-limiting. We saw that the general shape of repression-growth rate curves was similar for biochemically related genes – for example *dapA*/*dapB* (Fig 2E-F) or *purN*/*purL* (Fig 2J-K), in line with previous findings^11^.

In addition to these single gene relationships, we densely sampled pairwise expression-growth rate landscapes (Fig 3, lower triangle). In this matrix, every row and column represents a unique sgRNA, sorted by target gene and repression strength, and each pixel represents the growth rate effect of a given sgRNA pair. To quantify interactions between CRISPRi perturbations cooccurring within a cell, we quantified epistasis between every pair of CRISPRi knockdowns using a multiplicative (Bliss) model, where epistasis between sgRNAs 1 and 2 is equal to gr_1,2_ – gr_1_ * gr_2_ (Fig 3, upper triangle). For the majority of sgRNA pairs, we observed minimal epistasis. This is consistent with expectation given that: 1) many of the gene pairs are drawn from unrelated metabolic pathways and 2) many sgRNAs yielded only modest changes in gene expression. Across the library, however, we observed both positive and negative epistasis, particularly for sgRNAs with the strongest effects on gene expression. Our data also recapitulated the synthetic lethality previously reported for *gdhA* and *gltB* as negative epistasis^39^: we observed minimal growth rate defects when these genes were repressed individually, but severe growth rate defects when both genes were repressed together. However, even for gene pairs where epistasis was prevalent, the intensity of epistasis varied drastically within different expression regimes. Given how widely gene-gene epistasis varied based on the specific perturbations explored, we sought to discover a new way to summarize and model gene-gene epistasis across entire pairwise expression-growth rate landscapes.

**Figure 3.**
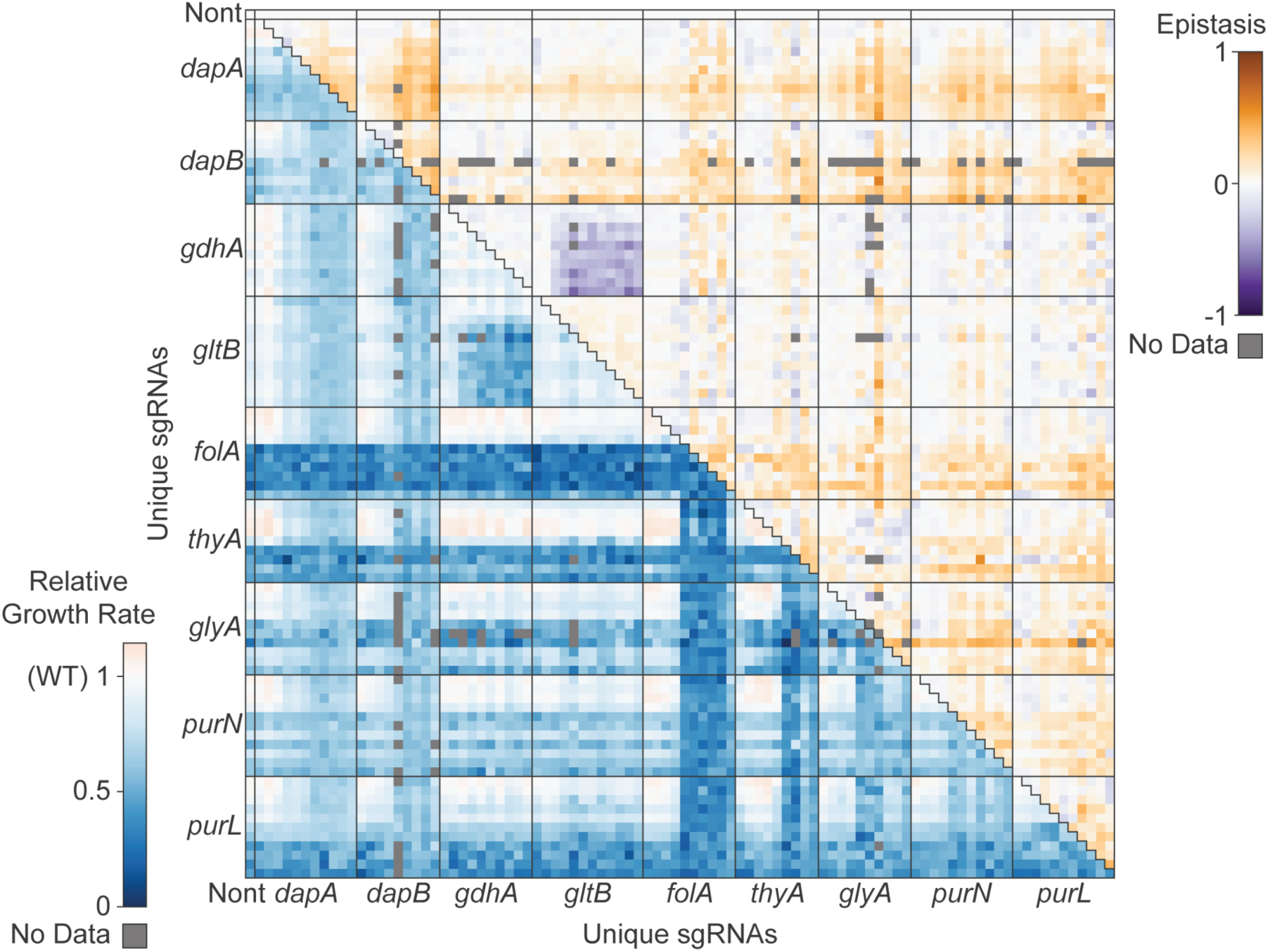
Pairwise CRISPRi growth rate and epistasis measurements. Each column and row represents a unique sgRNA perturbation. Gene names denote groups of sgRNAs targeting a given gene, and sgRNAs are sorted within each group by increasing CRISPRi repression strength (top-to-bottom and left-to-right). Nont indicates the non-targeting control sgRNA. The lower triangle of the matrix describes pairwise relative growth rate measurements calculated relative to the non-targeting control across 14 hours, averaged across n ≥ 4 experimental replicates, where wildtype-like (WT) growth is equal to 1. The upper triangle of the matrix describes pairwise growth rate epistasis. Epistasis was calculated as the difference between the observed growth rate and the multiplicative (Bliss) growth rate expectation of pairwise sgRNA perturbations.

### A continuous, coupling-sensitive model of expression-growth rate relationships

In developing the model, we sought to move beyond a matrix of many discrete epistasis measurements to a continuous, predictive description of expression-growth rate relationships that would be amendable to future studies of biological data. Under the assumption that densely measuring single perturbation-phenotype relationships is efficient (scaling linearly as the number of perturbations explored increases), we used well-constrained single perturbation functions as the backbone of our modeling strategy. To begin, we modeled the effect of single gene knockdowns on growth rate. We found that a sigmoidal model relating relative repression (R) to growth rate (g) using two parameters, steepness (n) and repression at half-maximal growth rate (R_0_) effectively modeled all single gene knockdown data (Equation 1, Fig 2E-M, Table S2).

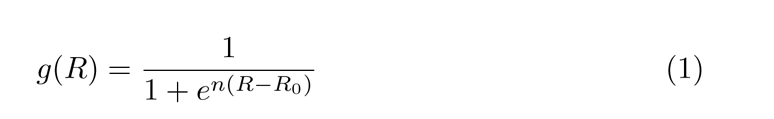

These sigmoidal fits enabled continuous prediction of the effects of gene repression on growth rate, improving upon our discrete experimental measurements. We used bootstrap resampling of the data to estimate 95% confidence intervals (CIs) for the sigmoidal fit and each parameter (Table S2, Methods). These fit parameters were well constrained for all genes except *gdhA* and *gltB* – two nonessential genes without significant growth rate defects. Although these two expression-growth rate functions can be fit by a wide range of n and R_0_ values, we will show that our model’s performance is insensitive to this variation and thus robust for essential and nonessential genes. We wanted to retain this continuous predictive power as we expanded our growth rate predictions to pairwise CRISPRi knockdowns. Assuming independence between CRISPRi perturbations, we constructed a Null model for pairwise knockdowns by multiplicatively combining the growth rate effects of two single gene knockdown functions (Equation 2).

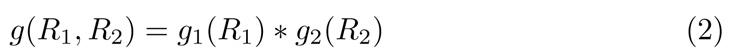

This initial model enabled continuous prediction of growth rate following pairwise gene knockdown but did not explicitly account for the possibility of gene-gene epistasis. To introduce epistasis into this modeling framework, we fit coupling constants a_ij_ and a_ji_ to modulate the effective repression of a given gene (R_ieff_) based on the intensity of a secondary perturbation (R_jeff_) and vice versa, as in Zimmer et al. 2016^28^ (Equation 3). These effective repression values are then used in place of true repression values in equations 1 & 2.

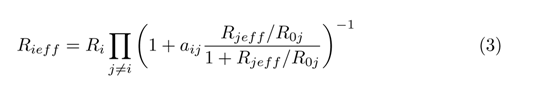

Based on the assumption of sparse gene-gene coupling, we introduced a regularization term to penalize the absolute values of a_ij_ and a_ji_ (Methods). Using mild regularization, we reduced the spread of coupling constant values while retaining our model’s predictive power (Fig S2, Table S3).

To investigate these pairwise expression-growth rate landscapes further, we highlight several example gene pairs. Firstly, for gene pair *dapA*/*purN* (Fig 4A), we observed that pairwise knockdowns (lower right pixels) had less severe growth rate defects than would be expected from single knockdown measurements alone (upper right and lower left). Our continuous epistasis model captured this interaction with a pair of positive gene-gene coupling terms (a_ij_ = 0.061, 95% CI [0.042, 0.099], a_ji_ = 0.197, [0.132, 0.238], Fig 4B), while the Null model erroneously underestimated growth (Fig 4C). Notably, the couplings between *dapA* and *purN* are asymmetric: in this instance, it reflects the fact that *purN* repression more strongly modulates the effect of *dapA* repression than the converse. In contrast, the synthetic lethal *gdhA* and *gltB* show negative epistasis, where the double knockdown phenotype was more severe than expected (Fig 4D). In our continuous epistasis model, this was captured with one negative gene-gene coupling term and one zero coupling term (a_ij_ = 0, [-1.06, 0], a_ji_ = -0.425, [-1.19, 0], Fig 4E), but this phenotype was again missed by the Null model (Fig 4F). In this case, the apparent coupling directionality for *gdhA*/*gltB* is the result of our regularization strategy and, as the confidence intervals indicate, either coupling constant can be negative to recapitulate synthetic lethality. The gene pair *dapA*/*dapB* shows positive coupling with near-symmetrical coupling constants, which we infer is because their gene products catalyze consecutive metabolic reactions (a_ij_ = 0.361, [0.237, 0.449], a_ji_ = 0.327, [0.267, 0.387], Fig S3A-C). Metabolic flux is limited by only a single knockdown, and the effect of an additional knockdown is dampened by this first flux restriction. However, we see almost no interaction between *purN* and *purL*, even though the products of these genes are also consecutive (a_ij_ = 0.053, [0.034, 0.095], a_ji_ = 0.009, [0.002, 0.016], Fig S3D-F). Thus, in the presence of symmetrical or asymmetrical coupling, the continuous epistasis model summarizes gene-gene coupling across entire pairwise expression-growth rate landscapes without incorporating bias from prior expectations.

**Figure 4.**
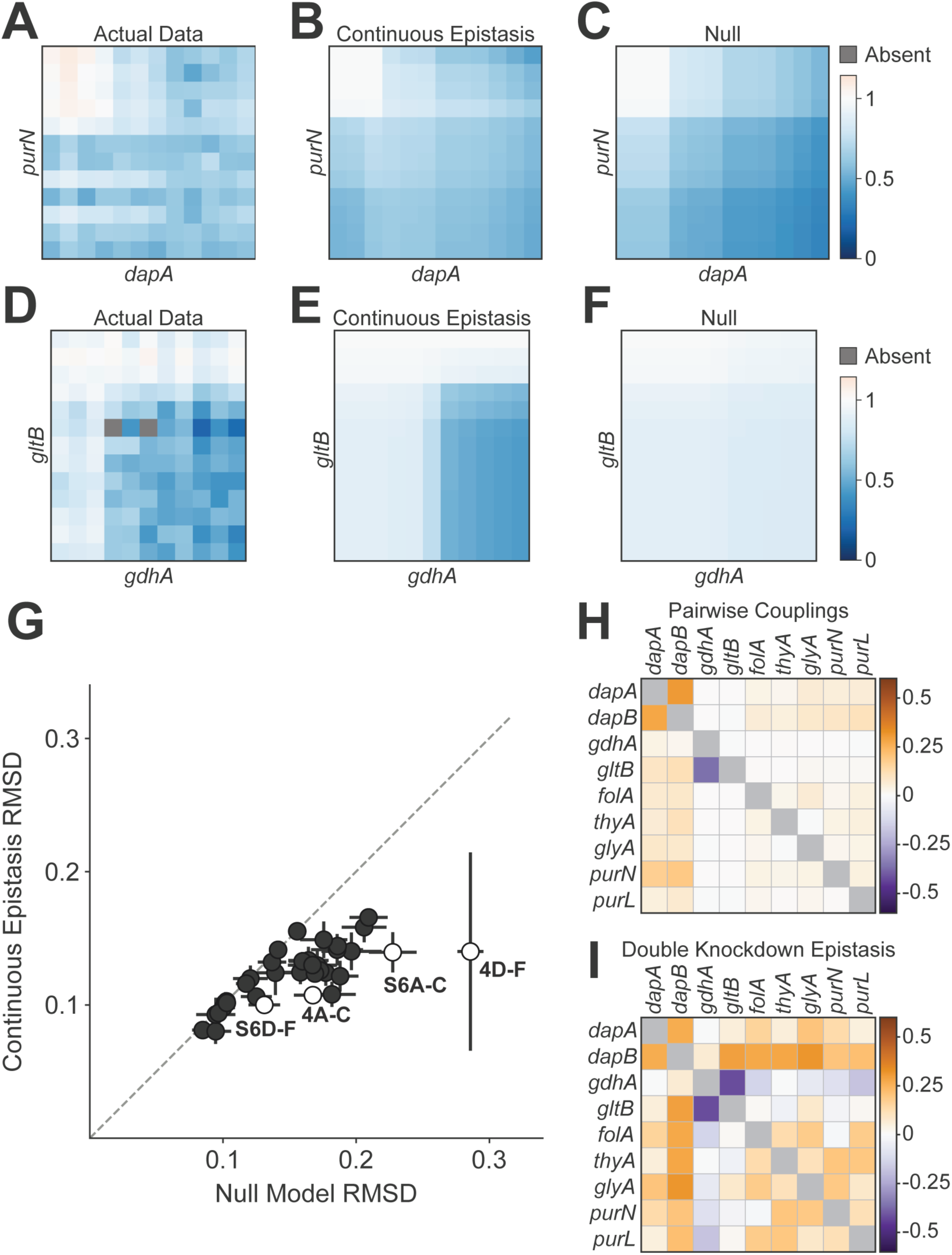
The continuous epistasis model captures gene-gene coupling at all expression levels. (A) Pairwise expression-growth rate data following CRISPRi knockdown of both *dapA* and *purN*, which show positive coupling. Each row and column represents a unique sgRNA, and pixels represent the growth rate effect of a given sgRNA pair. Rows and columns are sorted by knockdown intensity, ranging from wildtype-like expression (top left) to maximal double-knockdown (bottom right). (B) Predicted expression-growth rate data utilizing the coupling-sensitive continuous epistasis model. (C) Predicted expression-growth rate data utilizing the coupling-insensitive Null model. (D-F) Same as A-C, but for the *gdhA*/*gltB* gene pair, which shows negative coupling and synthetic lethality. (G) The performance of the coupling-sensitive model compared to the Null model. Each data point represents model RMSD across all pairwise sgRNA combinations for a gene pair. Full expression-growth rate data is shown for white, annotated dots in this figure and Fig S3. Error bars are standard deviations of models fit on bootstrapped single perturbation-growth curves. Dotted gray line is y = x. (H) Coupling constants fit across all pairwise growth rate data, as calculated from the continuous epistasis model. (I) Growth rate epistasis of the most severe double knockdown for each gene pair, calculated as the difference between the experimentally determined growth rate and the product of each single-gene knockdown growth rate.

The resulting epistatic model effectively predicted growth rates following knockdown for every gene pair, as quantified by the root-mean-square deviation (RSMD) between the model’s predictions and experimentally determined growth rates (Fig 4G). These decreases in RMSD (visualized as a point falling below y = x) illustrated that our model outperformed the nonepistatic Null model for most gene pairs. Indeed, when we calculated each model’s overall RMSD by pooling data from every gene pair together, our model (RMSD = 0.123, [0.122, 0.139]) significantly outperformed the Null model (RMSD = 0.158, [0.151, 0.173]). We wanted to ensure that these improvements were sufficient to warrant the additional complexity introduced by fitting coupling constants. Thus, we computed the Akaike Information Criterion (AIC) for each model, which supported the inclusion of these additional parameters (our model AIC = -15,716, [-15,785, -14,743], Null AIC = -13,958, [-14,299, -13,262]). We refer to this regularized, coupling-sensitive model as the “continuous epistasis” model.

Next, we sought to understand the relationship between the couplings learned by our approach and more standard couplings computed from “binary” genetic interaction measurements. We compared the gene-gene coupling values (a_ij_ and a_ji_) fit using our model to epistatic couplings calculated from a Bliss model considering only maximal-strength knockdown, mimicking a classical double-knockout study (Fig 4H-I). We found that coupling, when considered over an entire pairwise expression-growth rate landscape, is vastly less common than what is indicated by end-point perturbations alone. This insight highlights how seemingly disparate epistasis measurements can be concisely summarized when treated as a continuous landscape.

Finally, we tested two alterations to our model. For the first, we tried an alternate method with data requirements between the binary approach illustrated above and our continuous modeling technique by rigorously fitting single gene repression-growth rate curves, but fitting coupling constants using only the most extreme double-knockdown measurement. This allowed us to evaluate the utility of training on growth rate data following intermediate, pairwise knockdowns. This approach was prone to overfitting on the double-knockdown growth rate and failed to recreate intermediate phenotypes or accurately quantify epistasis (full model RMSD = 0.194, [0.174, 0.206]). As extreme double-knockdowns cause the most severe growth rate defects, and experimental noise in our system increases as growth rate decreases, this overfit model performs worse than the nonepistatic Null (Fig S3G). Second, to test whether the varying asymmetries we noticed in epistatic coupling constants were meaningful and indicative of true biological directionality, we transposed the coupling constants fit from our initial model, swapping a_ij_ and a_ji_ terms within each gene pair, and reassessed model performance. This transposed model performed significantly worse than the model with correct coupling directionalities, although it still outperformed our Null model as some epistatic coupling constants are truly symmetric (Fig S3H, RMSD = 0.137, [0.131, 0.154]).

### The expression-growth rate model can be reliably inferred using a small number of measurements

A key strength of this modeling strategy is its ability to represent an entire pairwise expression-growth rate landscape using a short list of parameters (n and R_0_ for each of two genes, a_ij_, and a_ji_). However, if the pairwise landscape must be densely sampled to reliably fit these parameters, a predictive model will be of little additional use. To quantify the minimal sampling requirements to accurately fit coupling constants, we evaluated the effect of subsampling our pairwise expression data and refitting the model on a smaller training set. As we wanted this analysis to inform library design strategies for future experiments, we tested three distinct subsampling patterns: subsampling a fraction of sgRNA pairs from across our entire library (Fig S4A), subsampling a fraction of sgRNA pairs for each gene pair in our library (Fig S4B), and subsampling a fraction of sgRNAs from our initial oligo pool, then constructing all pairwise combinations of the remaining sgRNAs (Fig S4C). The first approach would be implemented by constructing a large pairwise library and then bottlenecking prior to growth rate selection and sequencing, while the last is akin to this study with a smaller starting library. The middle approach captures an intermediate study (and may be more difficult to realize experimentally). Using each approach, we randomly split our pairwise expression-growth rate data into a training and test set, fit epistatic coupling constants for each gene pair using the training data, and evaluated the model’s growth rate predictions on our test data over 100 unique subsampling iterations. By varying the relative sizes of our training and test sets, we determined that the model is reliably fit with 20% of our original input data when subsampling using either of the first two strategies (Fig S4D-E). The third approach, subsampling sgRNAs from the starting library, was robust to subsampling down to 40% of the initial library size. Given these data, we recommend bottlenecking a diverse CRISPRi library as a simple but robust strategy to sparsely sample epistatic landscapes (RMSD = 0.128, [0.125, 0.144], Fig S5).

### High-order predictions can be made using low-order training data

We next wanted to evaluate whether a model trained on only single and pairwise expression-growth rate data could predict the effects of higher-order CRISPRi perturbations. This would vastly dampen the effect of the combinatorial explosion problem, enabling more complex expression-growth rate predictions. As such, we constructed a third-order sgRNA library and quantified growth rates following these third-order CRISPRi perturbations (Fig 5A). The library contained 911 distinct constructs consisting of 4 single, 76 pairwise, and 830 triple gene knockdowns and a non-targeting control (Fig 5B). Four additional non-targeting sgRNA controls were added to the library to ensure the presence of low-order sgRNA constructs in the final library. These alternative non-targeting sgRNAs phenocopy our original non-targeting control (Fig S6A). As each sequencing experiment measures growth rates relative to the overall culture and bulk culture doubling time varies based on library contribution, we rescaled relative growth rates from this experiment such that our single and pairwise CRISPRi knockdowns were well-correlated to those from our pairwise data set (RMSD = 0.118, Fig S6B).

**Figure 5.**
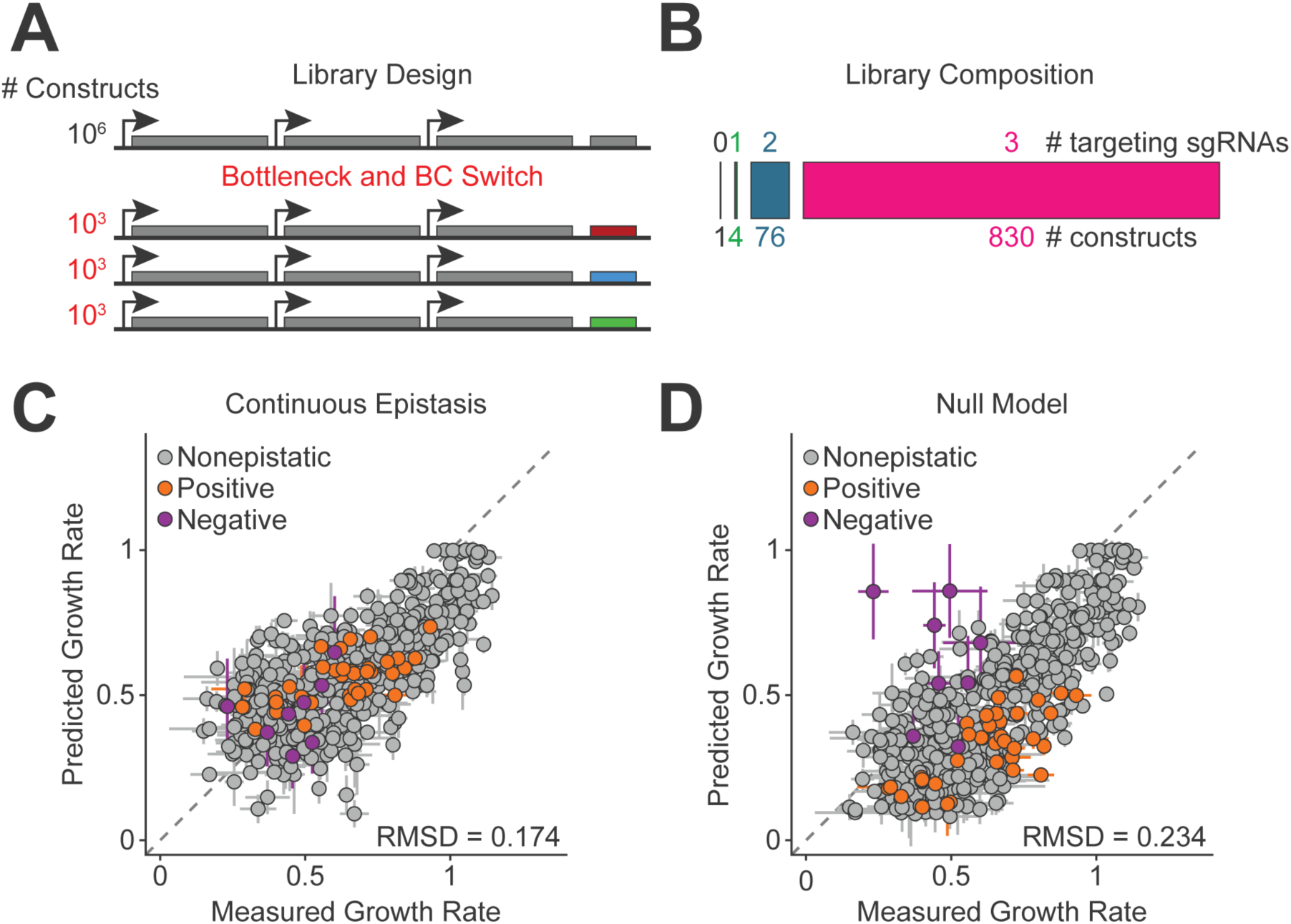
A model trained on pairwise measurements predicts growth rate following third-order gene knockdowns. (A) Diagram depicting the design of the third-order sgRNA library. After generating a diverse three-sgRNA library with >10^6^ possible constructs, we bottlenecked our library to a diversity of ∼10^3^ constructs. We then used inverse PCR to generate five copies of the library with identical sgRNA constructs but distinct barcodes (BC). (B) Distribution of targeting sgRNAs in the library. The library contained zero-through third-order targeting sgRNA combinations, with bars scaled to illustrate each order’s representation in the library. (C-D) Model performance on third-order sgRNA combinations, where experimentally determined growth rates are plotted against predicted growth rates for the (C) continuous epistasis and (D) Null model. Data is colored based on the presence of pairwise positive or negative epistasis (orange and purple points, respectively) within a construct. For a construct to be considered epistatic, two sgRNAs must target genes with a magnitude of epistasis (a_ij_ or a_ji_) > 0.15 with fewer than 9 sgRNA mismatches. Dotted gray line is y = x.

We used the gene-gene coupling values fit on our pairwise data to predict growth rates following all third-order CRISPRi perturbations. These predictions outperformed the coupling-insensitive Null model (RMSD = 0.174 vs RMSD = 0.234 for the epistatic and Null models, respectively, Fig 5C-D). Both models underestimate growth rate near wildtype, as these minor perturbations seem to combine sub-additively. However, when considering moderate and severe growth rate defects (where ∼75% of the data lie), the models’ predictions diverge. The continuous epistasis model recapitulates growth rates observed in our data, while the Null model continues underestimating growth. The accuracy of the Null model is also highly variable based on the presence or absence of epistasis between sgRNAs in a given construct, while the continuous epistasis model accounts for these interactions in its predictions and performs comparably across all epistatic regimes.

We then compared our model’s predictions to ones made using two alternative methods of estimating high-order phenotypes from low-order data: the Isserlis and regression-based models. These models lack interpretability, as they do not explicitly account for epistasis using learned parameters, and instead predict growth rates directly from single-and pairwise perturbation measurements. The models are also not inherently continuous, so we utilized “smoothed” versions of these models^28^. The Isserlis model performed comparably to our own, outperforming our model for growth rates near wildtype at the cost of increased uncertainty under slower growth conditions (RMSD = 0.160, Fig S6C). The regression-based model performed poorly due to its sensitivity to variance in low-order measurements (RMSD = 0.238, Fig S6D). Our model’s ability to extrapolate from pairwise data to predict the effects of higher-order gene knockdowns suggests that future studies of genetic epistasis across metabolic or transcriptional networks could be truncated at pairwise measurements, dampening the effect of combinatorial complexity.

### The continuous epistasis model captures gene-by-environment interactions

Importantly, the continuous epistasis model is unit-agnostic – because the model considers perturbation intensities on a relative scale, couplings need not be limited to gene-gene epistasis, but might more generally describe interactions between genes, environments, and other continuous perturbations. To assess the ability of the model to capture gene-by-environment interactions, we used mismatch CRISPRi to repress transcription of the folate metabolic genes *folA* and/or *thyA* while supplementing culture growth media with two products of folate metabolism: thymidine and/or methionine (Figure 6A, red text). Previous studies demonstrated that these perturbations have higher-order growth rate interactions^41–43^. In our own prior work, we observed that *folA* and *thyA* form an environmentally-dependent synthetic rescue. That is, growth rate defects due to loss of function mutations to DHFR (the gene encoded by *folA*) are rescued by loss of function mutations to TYMS (the gene encoding *thyA*) in the presence (but not the absence) of thymidine^41^. Moreover, the balance between thymidine starvation and amino acid limitation influences whether DHFR inhibition by the antibiotic trimethoprim results in growth inhibition or cell death^42, 43^. Thus, these combined genetic and environmental perturbations serve as an excellent test of our model’s ability to leverage pairwise couplings to predict high-order phenotypes.

**Figure 6.**
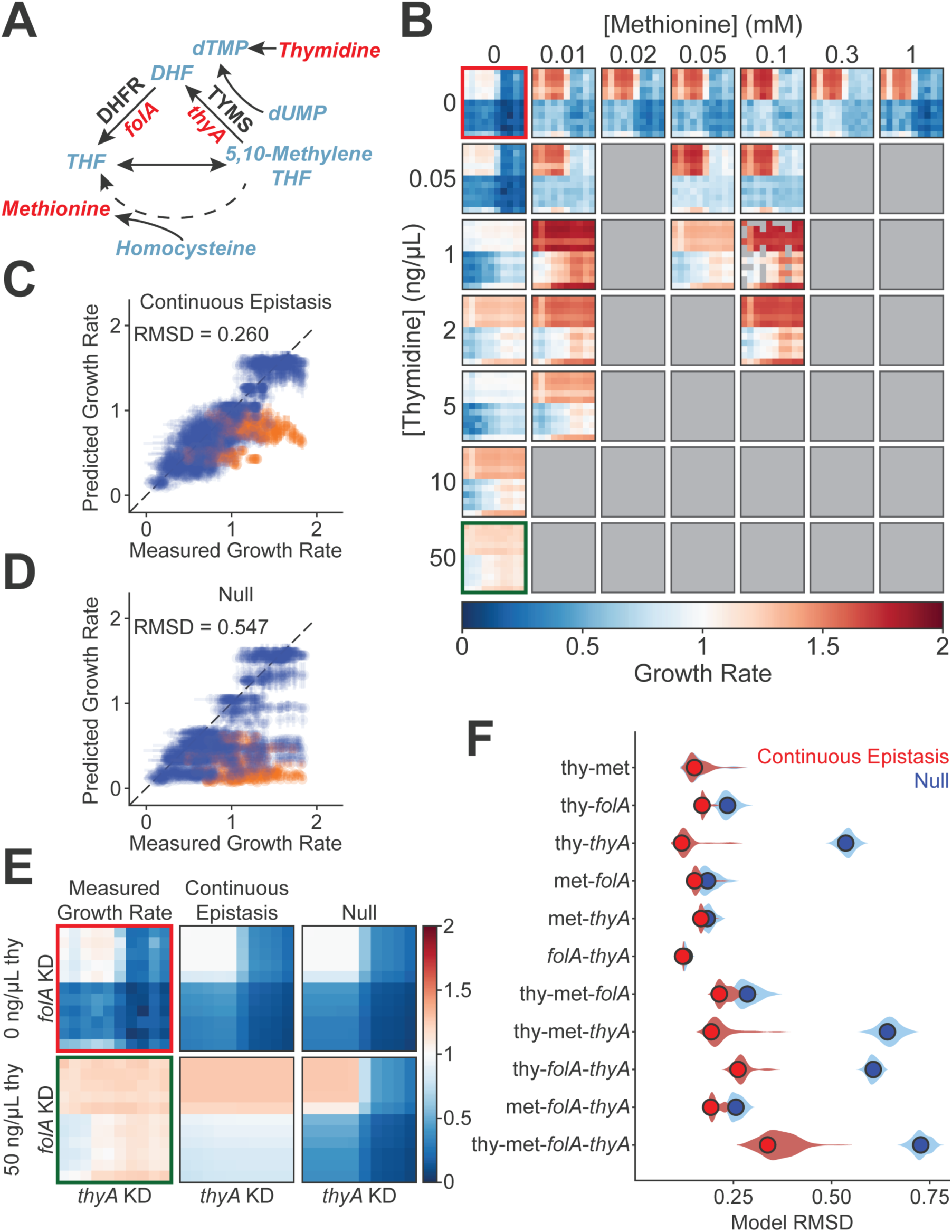
The continuous epistasis model captures gene-by-environment interactions. (A) Simplified schematic of a portion of *E. coli* folate metabolism. Gene names are listed in italics under each arrow, and enzyme abbreviations are listed in bold, black text above each arrow. Red text indicates a gene or metabolite perturbed over the course of the experiment. (B) Gene-by-gene-by-environment-by-environment relative growth rates, normalized to the growth rate of unsupplemented *E. coli* harboring a non-targeting control sgRNA. Within each heatmap, rows represent distinct *folA*-targeting sgRNAs, and columns represent distinct *thyA*-targeting sgRNAs. Thymidine and methionine supplementation varies between heat maps, and gray boxes represent pairwise media combinations not explored in this study. (C-D) Growth rate predictions for all data generated using the continuous epistasis (C) and Null model (D). Dotted line is y = x. The combined effects of *folA* knockdown, *thyA* knockdown, and thymidine supplementation result in a third-order interaction (orange points). (E) Expression-growth rate data and predictions following pairwise *folA*/*thyA* knockdown in unsupplemented media (top row) and media supplemented with 50 ng/μL thymidine (bottom row). (F) Continuous epistasis model performance, split by perturbation combination. For each second-and higher-order combination of expression and environmental perturbations (y-axis labels), the RMSD of growth rate measurements for the continuous epistasis model (red) and Null model (blue) are indicated. Violin plots represent model performance over 100 bootstrapped replicates.

To build the foundation of our gene-by-environment model, we first quantified single perturbation-growth curves by measuring the effect of media supplementation on bacterial growth rate in the absence of expression perturbations as well as the growth effect of expression perturbations in the absence of media supplementation. We measured growth rates for *E. coli* harboring a non-targeting CRISPRi control in a titrated range of thymidine and methionine concentrations. For wildtype *E. coli* in the absence of gene repression, the addition of either supplement to growth media slightly accelerated growth (Fig S7A-B, Table S4). These data were modeled with a 4-parameter sigmoid (using additional parameters g_min_ and g_max_ to represent the minimum and maximum growth rate effects, respectively) to quantify the growth rate effect of each environmental shift. We also refit repression-growth rate functions for *folA* and *thyA* using repression-growth rate data gathered in this experiment and bootstrapped all single perturbation-growth functions to estimate uncertainty in model performance and coupling strengths (Fig S7C-D, Table S5). To investigate gene-by-environment interactions within folate metabolism, we created a compact mismatch CRISPRi library containing 110 pairwise sgRNA combinations targeting *folA* and *thyA*. We transformed this library into our CRISPRi strain and grew the transformed population in a continuous culture device that allowed for parallel control of multiple supplemented media conditions^37, 44^. We again quantified relative growth rates following CRISPRi by next-generation sequencing and rescaled these measurements such that the growth rate of *E. coli* harboring the non-targeting CRISPRi control in unsupplemented media was equal to one. We then identified and removed escapers (125 measurements, 0.9% of data) and averaged across the remaining growth rate replicates, yielding 2,394 unique growth rate measurements across 22 growth conditions (Fig 6B). This generated a rich dataset to train and evaluate the continuous epistasis model on gene-by-environment interactions.

We isolated our pairwise (second-order) perturbation data and fit pairwise couplings between perturbations (Table S5). We found that our model effectively recapitulated growth rates for this independent data set, with an epistatic model RMSD of 0.145, [0.144, 0.176] and a Null RMSD of 0.273, [0.260, 0.293]. As with our gene-gene model, subsampling as little as 20% of our gene-by-environment data was sufficient to reliably fit the continuous epistasis model (RMSD = 0.151, [0.147, 0.194]). We performed subsampling using the same regularization strength used in our pairwise study, supporting the generalizability of this mild regularization. Thus, we see that our unit-agnostic model can capture not only gene-gene interactions, but also gene-by-environment and environment-by-environment interactions.

Next, we evaluated the ability of our model – trained exclusively on first-and second-order perturbations – to predict the growth rate effect of 2,035 third-and fourth-order perturbations. The continuous epistasis model outperformed the Null model on these higher-order predictions: including pairwise couplings yielded a prediction RMSD of 0.275, [0.252, 0.351] between the model and the data, while Null model predictions had an RMSD of 0.584, [0.555, 0.612]. High-order predictions remained robust when subsampling pairwise training data (RMSD = 0.300, [0.237, 0.412]). Combining all perturbations, the continuous epistasis model significantly outperformed the Null model on gene-by-environment predictions in terms of RMSD (continuous epistasis: 0.260, [0.239, 0.330], Null: 0.547, [0.521, 0.575]) and AIC (continuous epistasis: -6,416, [-6,817, -5,278], Null: -2,868, [-3,108, -2,637] (Fig 6C-D). We again compared our model to the Isserlis and regression models, and all models performed similarly (Isserlis RMSD = 0.300, regression RMSD = 0.254). However, only the continuous epistasis model yielded summarized coupling values between genes and environments in addition to growth rate predictions.

We can visualize why the continuous epistasis model so drastically outperformed the Null model using select environmental case-studies (Fig 6E). The top left matrix (red outline) contains growth rate measurements following CRISPRi knockdown in unsupplemented M9-glucose media, and as expected, knockdown of either gene is deleterious to growth. Both the continuous epistasis and Null models accurately recapitulated this experimental data as there is not strong pairwise epistasis between *folA* and *thyA* in this environment. However, upon addition of thymidine into the media (bottom left, green outline), the models’ predictions diverge. The addition of thymidine to the media decouples *thyA* expression from growth rate. We observe this phenotype in the first few rows of these matrices, where thymidine knockdown does not cause a growth defect. This phenotype is captured by the continuous epistasis model, but the Null model’s predictions do not change following nutrient supplementation except to uniformly increase growth rate in the richer media condition. We also note that a phenotype present in our experimental data is missing from both the continuous epistasis and Null models – when thymidine is present in the media and *folA* is strongly repressed, *thyA* repression actually *benefits* growth. As mentioned above, this third-order gene-by-gene-by-environment interaction was expected and is not captured by our pairwise model (Fig 6C-D, orange points). This highlights the need for careful evaluation of model performance, as a systematic failure in a pairwise model may be indicative of biologically interesting high-order interactions. Lastly, we observed a “stripe” of slightly elevated growth in the last row of our pairwise expression-growth rate matrices. This apparent nonmonotonicity likely arises because matrix rows are ordered by sgRNA repression strength, and the strongest knockdowns (bottommost rows) are nearly indistinguishable by qPCR and thus may be misordered. This sensitivity highlights the importance of the bootstrapping approach we used when fitting single perturbation-growth functions, as we know the model’s overall performance is generally insensitive to these variations at maximal knockdown.

To quantitatively, rather than qualitatively, investigate where our model’s accuracy gains emerged, we separated our data by genetic and environmental perturbation types and plotted each model’s error separately for each perturbation subset (Fig 6F). The continuous epistasis model outperformed the Null model the most when *thyA* was repressed and thymidine was added to the media. As noted previously, thymidine replaces the key metabolite made by the *thyA* gene product, rescuing growth and resulting in a positive pairwise coupling (a_ij_ = 0, [0, 0], a_ji_ = 0.73, [0.20, 1.29]). This asymmetric coupling is indicative of a hierarchy between thymidine and *thyA* expression, where the addition of thymidine to growth media fully decouples growth rate from *thyA* expression but not vice versa. These results indicate that the continuous epistasis model can be applied beyond low-order perturbation studies to predict the combined effects of complex gene-by-environment and environment-by-environment interactions. This now opens the door to creating continuous, coupling sensitive pharmacogenomic models that describe the interaction between drug, genotype, and environment.

## Discussion

In this work, we generated and trained a continuous, coupling-sensitive model relating gene expression to growth rate using sparsely-sampled single and pairwise perturbation measurements. The model’s parameters are interpretable and relevant across gene expression levels, and this unitless approach enables the investigation of context-dependence between disparate perturbations, such as CRISPRi knockdown and metabolite supplementation. We observed that gene-gene epistasis, when inferred across a continuous expression landscape, is less common than epistasis inferred at the level of knockout or extreme knockdown. In addition, while it is possible to infer why some gene pairs show positive or negative coupling based on prior knowledge, we see that biochemical pathway structure is not sufficient to predict all gene-gene interactions, and it does not seemingly offer perspective on coupling strength. Certainly, the relationship between coupling strength, asymmetry, and functional interactions between enzymes will be an interesting avenue for further study.

Based on our results, we propose a hybrid experimental/modeling approach to streamline epistatic measurements, generate quantitative insight into the strength and directionality of epistasis, and predict the outcome of perturbations not explored within a given study. First, precisely quantify the effect of each single perturbation at varied intensities (Fig 7A). This can be accomplished for gene knockdown measurements using mismatch CRISPRi or for drug treatment by sampling around a known IC_50_. Then, sparsely sample pairwise perturbation landscapes (exact sampling density may vary; this study used 10s of measurements per combination) (Fig 7B). Using the continuous epistasis model, fit pairwise couplings to quantify the magnitude and directionality of pairwise epistasis between all perturbation combinations (Fig 7C). Finally, utilize these epistatic couplings and single perturbation fits to continuously model each pairwise perturbation landscape and predict the effect of high-order perturbation combinations (Fig 7D). For reference, if one wished to perform an experiment with the same sampling density (20 measurements per pair), replicates (6), and sequencing depth (≥70x coverage) as the study discussed here – but for 1,000 genes of interest (totaling ∼1 million epistatic interactions and half a million pairwise landscapes) – the sequencing data required would fit on a single NovaSeq X 25B flow cell.

**Figure 7.**
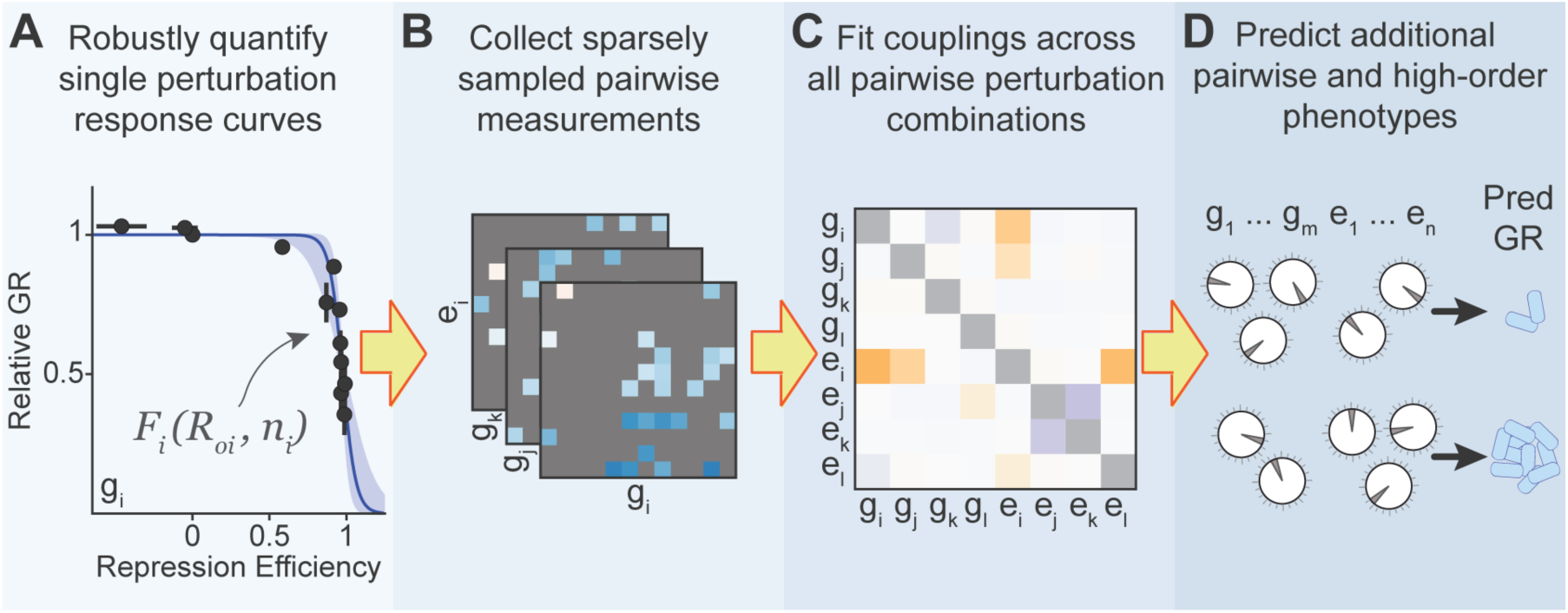
An overview of epistasis studies using the continuous epistasis model. (A) Densely sample single perturbation response functions using individual measurements, quantifying R_0_ and n for each individual perturbation. (B) Collect sparse pairwise measurements between all individual perturbations, randomly sampling the pairwise landscape. g_i_: gene i; e_i_: environment i. (C) Use the single perturbation response functions and sparse pairwise data to fit pairwise couplings between all perturbation combinations. (D) Use the continuous epistasis model to model continuous single, pairwise, and higher-order landscapes and predict phenotypes for unmeasured perturbation combinations.

While we limited our experiments to transcriptional and environmental perturbations in this work, one can imagine integrating proteomic, metabolomic, and other measurements into this unitless modeling framework. We modeled the growth rate effect of each perturbation with a monotonic function, however there are known examples of nonmonotonic growth rate relationships, limiting some applications of the model as it is currently constructed^10^. Finally, while we showed that accounting for pairwise coupling alone captured most variation in our third-and fourth-order perturbation landscapes, sufficiently strong high-order epistasis in a system could require a different approach.

To expand this workflow, the limiting experimental step is quantifying the repression strength of individual sgRNAs. RT-qPCR is relatively low-throughput, precluding genome-level studies. An alternative, single-cell approach to quantify CRISPRi repression, such as microSPLiT^45^ (bacterial cells), PETRI-seq^46^ (bacterial cells) or perturb-seq^47^ (mammalian cells) alongside next-generation sequencing-based growth rate measurements would remove these throughput restrictions and permit significantly larger CRISPRi studies. Alternatively, one could utilize a predictive model relating sgRNA homology sequence to CRISPRi repression strength^11, 32^.

Applications of this continuous, epistatic model are numerous. The reduced sampling requirements make pathway-level epistatic analyses feasible in single experiments. Prior work has indicated that metabolic enzymes are often produced in evolutionarily conserved stoichiometric ratios^48^, and the experiments presented here could be used to better understand the constraints on enzyme abundance in metabolic systems that shape these ratios. The continuous epistasis model could also be applied to optimize biosynthetic pathway performance and/or cell growth conditions. We hope that interpretable, continuous, and coupling-sensitive models like the one described in this work can continue to improve computational predictions to extract more information from the data-rich -omics experiments we now regularly perform.

## Acknowledgments

The authors would like to thank Olivier Rivoire for insightful discussions and data analysis ideas. We are grateful to Prashant Mishra for thoughtful comments on our manuscript. We thank Andrew Mathis for feedback in the early stages of this work. We also thank the Reynolds lab for feedback on the project and manuscript. Research reported in this publication was supported by the National Institute of General Medical Sciences of the National Institutes of Health under award number R01GM136842. R.O. was also partly supported by a National Institutes of Health training grant 5T32GM007062-46. This content is solely the responsibility of the authors and does not necessarily represent the official views of the National Institutes of Health.

## Author Contributions

Conceptualization, R.M.O. and K.A.R.; Formal Analysis – R.M.O., A.T.N., P.M.B., and K.A.R.;

Investigation – R.M.O., A.T.N., and P.M.B.; Writing – Original Draft, R.M.O., P.M.B., and K.A.R.; Writing – Review & Editing, R.M.O., A.T.N., P.M.B., and K.A.R.; Supervision, K.A.R.; Funding Acquisition, K.A.R.

## Methods

### Pairwise CRISPRi Library Construction

We designed a pool of CRISPRi sgRNAs targeting nine *E. coli* K-12 MG1655 genes. For each gene, we selected 9-12 distinct sgRNAs to provide a titrated range of mRNA knockdown. We previously characterized CRISPRi knockdown of seven of these genes^33^ (*dapA*, *dapB*, *folA*, *thyA*, *glyA*, *purN*, and *purL*), giving us an initial list of sgRNAs for each gene with mutually distinguishable growth rate effects. We supplemented these sgRNAs with up to two fully on-target sgRNAs per gene and added sgRNAs by hand to populate underrepresented growth regimes. The final two genes (*gdhA* and *gltB*) were previously uncharacterized by our lab. We designed two on-target sgRNAs for each gene using previously published design principles^30^. In addition, we introduced 4, 6, 8, 10, and 14 mismatches into the homology region of each on-target sgRNA^33^. For a full list of sgRNA sequences, see Table S1.

All sgRNAs were synthesized as a mixed oligo pool by Integrated DNA Technologies. We resuspended the library to a final concentration of 50 nM in MB H_2_O and added BsaI flanking sites using PCR. We performed two separate PCRs to add two distinct sets of flanking sites onto the library for Golden Gate cloning. We used the GG_UnivF/GG_R1 primers to generate the insert 1 library and the GG_F2/GG_UnivR primers to generate the insert 2 library (for primer sequences, see Table S6). The PCR mix contained 10.75 μL MB H_2_O, 5 μL Q5 Reaction Buffer (NEB), 5 μL High GC Enhancer (NEB), 0.5 μL 10 mM dNTPs, 1.25 μL 10 μM forward primer, 1.25 μL 10 μM reverse primer, 0.25 μL Q5 High-Fidelity DNA Polymerase (NEB, #M0491), and 1 μL of the 50 nM oligo pool. Initial denaturation was carried out at 98°C for 30 seconds, followed by 25 cycles of 98°C for 30 seconds, 58°C for 10 seconds, and 72°C for 20 seconds. A final elongation was performed at 72°C for 60 seconds, followed by an indefinite hold at 4°C. We ran the PCR product on a 1% agarose 1X TAE gel with EtBr staining at 80 V for 45 minutes and excised the 176 bp band. We extracted the band using a DNA Clean & Concentrator-5 Kit (Zymo, #D4014), eluted in 30 μL MB H_2_O, and quantified dsDNA concentration using a PicoGreen assay (Thermo, #P7581). To ensure representation of the non-targeting sgRNA sequence (Nont) in the final library, we isolated this control sgRNA using the same protocol illustrated above, replacing the 50 nM oligo pool with 1 μL of purified 20-50 ng/μL plasmid containing the Nont sgRNA^33^ (pCRISPR2). We spiked Nont inserts into each of the mixed libraries at a 1:100 Nont:library ratio.

We performed three Golden Gate cloning reactions to ligate the CRISPRi inserts into three distinct barcoded CRISPRi vectors (pCRISPR3, AddGene #: 191856-191861). The reaction mix contained 75 ng of barcoded vector, 6 ng of insert 1 mix, 6 ng of insert 2 mix, 1.5 μL of 10X T4 DNA Ligase Buffer (NEB), 1 μL T4 DNA Ligase (2,000,000 U/mL, NEB, #M0202M), 1 μL BsaI-HF v2 (NEB, #R3733), and MB H_2_O to 15 μL. Reaction conditions were 40 cycles of 37°C for 2 minutes and 16°C for 3 minutes. These cycles were followed by 50°C for 10 minutes, 80°C for 20 minutes, and an indefinite 4°C hold. A no-insert control, where both inserts were left out of the reaction, was performed identically.

We cleaned and concentrated each reaction using a DNA Clean & Concentrator-5 Kit (Zymo, #D4014), eluting in 20 μL MB H_2_O. We transformed 2 μL of each purified library into 100 μL of XL1 Blue electrocompetent cells followed by a 1-hour recovery at 37°C with shaking. We plated 50 μL of a 1:10 and 1:100 dilution to quantify library coverage and added the remaining cells to 9 mL SOB + 35 μg/mL kanamycin. We incubated this culture overnight at 37°C with shaking. We also transformed and plated the no-insert control and a no DNA control and adjusted library coverage estimates accordingly. Library coverage ranged from 10-14x for each barcoded vector. In addition to these mixed CRISPRi libraries, we synthesized constructs containing two Nont sgRNAs in six distinct barcoded vectors using the same protocol described above, substituting each of the mixed insert libraries for 6 ng of purified Nont sgRNA insert.

We isolated plasmids using the GeneJET Plasmid Miniprep Kit (Thermo, #K0503), eluted DNA in 50 μL MB H_2_O, and quantified dsDNA concentration with a PicoGreen assay (Thermo, #P7581). We combined the three barcoded CRISPRi libraries in an equimolar ratio. We also mixed the Nont+Nont controls in an equimolar ratio, then spiked this control into the library at a 1:5,000 Nont+Nont:library ratio. We transformed 3 ng of this final mixture into 45 μL MG1655 + chromosomal dCas9 electrocompetent cells (gift from the Bikard lab) using the same transformation protocol as above, plating 50 μL of 1:100 and 1:1,000 dilutions. We performed a no DNA control with zero breakthrough colonies. Coverage exceeded 200x, and we grew the remaining cells in 9 mL SOB + 35 μg/mL kanamycin overnight at 37°C with shaking.

The pairwise *folA*/*thyA* CRISPRi library was constructed as described above, with only Nont, *folA*-targeting, and *thyA*-targeting sgRNAs included in the Integrated DNA Technologies-synthesized oligo pool. Library coverage calculated after the first transformation event in the three barcoded vectors ranged from 14x to 89x. We mixed three barcoded Nont+Nont CRISPRi constructs in equimolar ratios and spiked them into the *folA/thyA* library at a 1:400 Nont+Nont:library ratio. As two continuous culture experiments were required to investigate all environments of interest, we freshly transformed the library two separate times. For each transformation, we added 1 ng of mixed library to 45 μL MG1655 + dCas9 electrocompetent cells and achieved 583x and 127x library coverage.

### Continuous culture

Following transformation of the MG1655 dCas9 electrocompetent *E. coli*, cells were grown overnight in LB + 35 μg/mL kanamycin. We spun saturated outgrowth cultures at 1,000 rcf for 5 minutes, decanted media, and resuspended cells in 1 mL M9 minimal media pH 6.5, 0.4% glucose, 35 μg/mL kanamycin (M9+Kan). We repeated this wash, then resuspended 10 μL of concentrated cells in 8 mL M9+Kan. We incubated this culture for 12 hours at 37°C with shaking.

Following outgrowth, we spun the culture at 1,000 rcf for 5 minutes, decanted, and resuspended cells in 1 mL fresh M9+Kan. We repeated this wash, diluted the culture to OD_600_ = 0.05 in 15 mL M9+Kan, and grew these cells in a continuous culture device at 37°C, clamping the optical density at 0.15. Our continuous culture device follows the design specifications from Toprak et al., 2012 and 2013^37, 44^. Following an overnight adaptation, we added anhydrotetracycline (Cayman Chemical Company, #10009542) to a final concentration of 50 ng/mL to induce CRISPRi. We took the first timepoint (T_0_) after 3 hours of CRISPRi by extracting 1 mL of cells from the culture and took additional timepoints every 2 hours until T_14_. We centrifuged timepoints at 3,000 rcf for 5 minutes to pellet cells, decanted by pipetting, and stored them at −20°C.

We performed continuous culture for the third-order CRISPRi and environmental variation experiments with slight modifications. We terminated these experiments after T_10_, as the final two timepoints were shown to be unnecessary for growth rate quantification. We inoculated the third-order CRISPRi library into the continuous culture device at a starting OD_600_ = 0.2. Media conditions were altered during the *folA*/*thyA* gene-by-environment experiment. Thymidine (Sigma, #T1895) and/or methionine (Sigma, #M5308) were supplemented into M9+Kan to create the following conditions: No additive, 0.05 ng/μL thymidine, 1 ng/μL thymidine, 2 ng/μL thymidine, 5 ng/μL thymidine, 10 ng/μL thymidine, 50 ng/μL thymidine, 0.01 mM methionine, 0.02 mM methionine, 0.05 mM methionine, 0.1 mM methionine, 0.3 mM methionine, 1 mM methionine, 0.05 ng/μL thymidine + 0.01 mM methionine, 0.05 ng/μL thymidine + 0.05 mM methionine, 0.05 ng/μL thymidine + 0.1 mM methionine, 1 ng/μL thymidine + 0.01 mM methionine, 1 ng/μL thymidine + 0.05 mM methionine, 1 ng/μL thymidine + 0.1 mM methionine, 2 ng/μL thymidine + 0.01 mM methionine, 2 ng/μL thymidine + 0.1 mM, and 5 ng/μL thymidine + 0.01 mM methionine.

### Sample preparation and next-generation sequencing

We resuspended each frozen cell pellet (each corresponding to a sample – one time point and environmental condition) in 100 μL MB H_2_O with vortexing and lysed them at 95°C for 3 minutes. Following a 10 minute, 20,000 rcf centrifugation to pellet debris, we took 50 μL of the supernatant as a working stock for use in downstream reactions.

We then added adaptors for Illumina sequencing in two consecutive PCR steps. First, we added flanking sequences to each timepoint or environmental sample. The reaction mix was composed of 10.5 μL MB H_2_O, 5 μL Q5 Reaction Buffer (NEB), 5 μL 50% glycerol, 0.5 μL 10 mM dNTPs, 1.25 μL 10 μM ΤruSeqF primer, 1.25 μL 10 μM TruSeqR_trunc primer, 0.5 μL Q5 High-Fidelity DNA Polymerase (NEB, #M0491), and 1 μL timepoint working stock. Reaction conditions were 98°C for 30 seconds, 7 cycles of 98°C for 30 seconds, 61°C for 10 seconds, and 72°C for 15 seconds, followed by a final elongation at 72°C for 45 seconds and an indefinite hold at 4°C. In the second PCR, we added Illumina sequencing adapters and i5/i7 barcoding sequences to each sample. The reaction mix contained 10.5 μL MB H_2_O, 5 μL Q5 Reaction Buffer (NEB), 5 μL 50% glycerol, 0.5 μL 10 mM dNTPs, 1.25 μL 10 μM i5 primer, 1.25 μL 10 μM i7 primer, 0.5 μL Q5 High-Fidelity DNA Polymerase (NEB, #M0491), and 1 μL template from the previous reaction. Reaction conditions were 98°C for 30 seconds, 20 cycles of 98°C for 30 seconds, 61°C for 10 seconds, and 72°C for 20 seconds, followed by a final elongation at 72°C for 60 seconds and an indefinite hold at 4°C.

Note: While the glycerol was used in the preparation of timepoints for the pairwise CRISPRi library experiment and the *folA*/*thyA* sublibrary experiment, we have observed future amplification reactions that failed in the presence of glycerol.

We quantified dsDNA concentration from each reaction using a PicoGreen assay (Thermo, #P7581), and mixed timepoints in an equimolar ratio to create the mixed sequencing library. We ran the library on a 1% agarose 1X TAE gel with EtBr staining at 90 V for 60 minutes and excised the 486 bp construct. We performed gel extraction using the DNA Clean & Concentrator-5 Kit (Zymo, #D4014) and eluted in 20 μL MB H_2_O. Following a Qubit Assay (Thermo, #Q32851) to quantify final dsDNA quantification, we submitted the library to Azenta for Illumina HiSeq Sequencing on a 300-cycle paired-end run.

### Isolation of single CRISPRi knockdown strains

To quantify the concentration of mRNA following each individual CRISPRi perturbation, we isolated plasmids expressing a single targeting sgRNA paired with the Nont sgRNA. For this purpose, we constructed a CRISPRi:Nont library using Golden Gate cloning as described previously, adding the Nont sgRNA insert in place of the mixed library in the second sgRNA position. We screened individual colonies from this mixed library to isolate the majority of CRISPRi:Nont constructs.

We individually synthesized plasmids not isolated during this screen using inverse PCR (iPCR). First, we phosphorylated iPCR primers with a PNK reaction. The reaction mix consisted of 6.5 μL MB H_2_O, 1 μL 10X T4 DNA Ligase Buffer (NEB), 1 μL 100 μM iPCR_sgRNA_F primer, 1 μL 100 μM iPCR_UnivR primer, and 0.5 μL T4 PNK (NEB, #M0201S). Thermocycler conditions were 37°C for 30 minutes, 65°C for 20 minutes, and an indefinite 4°C hold. Then, we performed iPCR with a reaction mix of 8.25 μL MB H_2_O, 5 μL Q5 Reaction Buffer (NEB), 0.5 μL 10 mM dNTPs, 0.5 μL Q5 High-Fidelity DNA Polymerase (NEB, #M0491), 10 μL primer phosphorylation reaction, and 1 μL 30-50 ng/μL template plasmid (pCRISPR2). Reaction conditions consisted of 98°C for 30 seconds, followed by 25 cycles of 98°C for 30 seconds, 58°C for 30 seconds, and 72°C for 90 seconds. A final 3-minute elongation at 72°C preceded an indefinite hold at 4°C. We cleaned and concentrated the reaction product using a DNA Clean & Concentrator-5 Kit (Zymo, #D4014) and eluted in 25 μL MB H_2_O. We removed the template plasmid using a DpnI digestion. Reaction conditions were 13 μL cleaned and concentrated product, 2.5 μL 10x Cutsmart Buffer (NEB, #B6004), 8.25 μL MB H_2_O, and 1.25 μL DpnI (NEB, #R0176). The reaction was held at 37°C for 1 hour, then held at 4°C. Finally, we circularized the PCR product in a ligation reaction consisting of 8 μL DpnI product, 1 μL 10X T4 DNA Ligase Buffer (NEB), and 1 μL T4 DNA Ligase (400,000 U/mL, NEB, #M0202S). Ligation occurred at 25°C for 2 hours, followed by 10 minutes at 65°C and a hold at 4°C. We transformed this product into XL1 Blue chemically competent cells (1 μL DNA, 45 μL competent cells) with 1 hour recovery at 37°C with shaking. We purified plasmids using a GeneJET Plasmid Miniprep Kit (Thermo, #K0503) and transformed into CRISPRi MG1655 + dCas9 cells as described previously.

### RNA Extraction and RT-qPCR

To collect RT-qPCR data following CRISPRi, we grew CRISPRi-Nont strains from glycerol stocks in 4 mL LB + 35 μg/mL kanamycin for 6-8 hours at 37°C with shaking. We washed these cultures into M9+Kan as described previously and incubated them for 8-14 hours at 37°C with shaking. We then washed cultures into M9+Kan+50 ng/mL anhydrotetracycline (Cayman Chemical Company, #10009542) and incubated them for 5 hours at 37°C with shaking at a starting OD_600_ = 0.05. After treatment, we isolated RNA using the Qiagen RNeasy Protect Bacteria Mini Kit (QIAGEN, #74524), with on-column DNase digestion (QIAGEN, #79254), as described in their protocol. We extracted RNA from control Nont+Nont strains using the same procedure. We performed RT-qPCR on a CFX Opus 384 Real-Time PCR System (Bio-Rad, #12011452) using the Luna Universal One-Step RT-qPCR Kit (NEB, #E3005). No template and genomic DNA controls were included in all experiments. We analyzed raw results using the CFX Maestro Software (Bio-Rad), and calculated changes in mRNA concentration using the ΔΔC_t_ method with the *hcaT* gene as reference in a custom Python script.

### Third-order sgRNA library construction and next-generation sequencing

We constructed the third-order sgRNA library in a single barcoded vector (pCRISPR3, AddGene #: 191856-191861) using Golden Gate cloning as described previously with minor modifications. Three sgRNA inserts were used instead of two, using primers GG_UnivF/GG_R1 for insert 1, GG_F2/GG_R2 for inserts 2, and GG_F3/GG_UnivR for insert 3 (Table S6). During Golden Gate cloning, insert concentrations were doubled to 12 ng per insert per reaction, and reaction conditions were adjusted to 60 cycles of 2 minutes at 37°C and 3 minutes at 16°C. These cycles were followed by 10 minutes at 50°C, 20 minutes at 80°C, and an indefinite 4°C hold. Transformed cells were serially diluted and plated, and we selected a library dilution containing ∼1,000 colony forming units to form our bottlenecked library.

This library was used as a template in an iPCR reaction to replace the 6-nucleotide barcode with a new barcode sequence while maintaining sgRNA diversity. iPCR was carried out as described above using primers BC_iPCR_F_BC[1-6] and BC_iPCR_UnivR_v2. These libraries were sequence verified and transformed at sufficient efficiency to avoid further bottlenecking. These sublibraries were quantified by Qubit and combined in an equimolar ratio to generate the final third-order CRISPRi library with five barcodes. pCRISPR3 barcode 4 was excluded from the reaction. Finally, to ensure that we could normalize sgRNA constructs to a wildtype-like subpopulation, CRISPRi-sensitive *E. coli* harboring a Nont+Nont+Nont construct (in all five barcodes) were spiked into the final population at a 1:500 ratio before CRISPRi treatment.

We altered the sampled preparation protocol to accommodate the increased amplicon size of the third-order sgRNA CRISPRi library. For the first PCR, we used a reverse primer with 4 additional base pairs of homology (TruSeqR). The reaction conditions were changed to 30 seconds at 98°C, 17 cycles of 30 seconds at 98°C and 60 seconds at 76°C, followed by 180 seconds at 72°C and an indefinite 4°C hold. In the second PCR, we altered thermal cycler conditions to 30 seconds at 98°C, 10 cycles of 30 seconds at 98°C and 75 seconds at 72°C, followed by 180 seconds at 72°C and an indefinite 4°C hold. We quantified dsDNA from these reactions using a Qubit Assay (Thermo, #Q32851), mixed them in an equimolar ratio, and concentrated DNA using a DNA Clean & Concentrator-5 Kit (Zymo, #D4014) with a 30 μL MB H_2_O elution. We ran this concentrated library on a 2% agarose 1X TAE gel with an EtBr stain at 90 V for 85 minutes and extracted the resulting 636 bp band using a DNA Clean & Concentrator-5 Kit (Zymo, #D4014) with a 20 μL MB H_2_O elution. We quantified dsDNA with a Qubit Assay (Thermo, #Q32851) and sequenced in-house using two Illumina MiSeq Nano v2 500-cycle paired-end runs and one Illumina MiSeq v2 500-cycle paired-end run.

### Plate reader growth rate assay

We cultured *E. coli* strains harboring a non-targeting CRISPRi control overnight in 4 mL LB + 35 μg/mL kanamycin at 37°C with shaking. The next day, we washed these cultures twice into M9+Kan as described previously. We diluted cultures 1:200 into each of the 22 media conditions described in Continuous Culture and adapted cells for 4 hours at 37°C with shaking. Following adaptation, we washed cells into 4 mL of their respective supplemented media + 50 ng/mL anhydrotetracycline at OD_600_ = 0.005. Three 200 μL replicates of each culture were grown in a Synergy Neo2 Hybrid Multi Mode Reader (Agilent, #BTNEO2) overnight at 37°C while taking regular OD_600_ measurements. Growth rates were fit across replicates using a log-linear fit of exponential phase growth and averaged across triplicate measurements.

### Calculating CRISPRi growth rates

We wrote all analysis code in Python 3.9.12 using Jupyter Notebook (available on github at https://github.com/reynoldsk/ContinuousEpistasis). We extracted sgRNA sequences from Illumina FASTQ files using regular expressions, searching for a 20 bp region between flanking sequences CTAGCTCTAAAAC and A. Up to 3 mismatches and no gaps were permitted in the flanking sequences. The identified homology region was required to match a desired sequence from the CRISPRi library. In addition, to increase confidence, we imposed a filter requiring two specific base calls in each sequence to have a Q-score above 30. The filtered bases were the final “on-target” and first “mismatch” base in the homology region, as this two-nucleotide combination is unique to a single sgRNA within each sgRNA family. We identified the plasmid barcode similarly, with flanking sequences GTACAGCGAGGCAAC and ACGGATCCCCAC, imposing a maximum of 3 mismatches and allowing no gaps. The barcode sequence needed to match a target barcode exactly, and no Q-score filter was imposed.

For the three-sgRNA library, the following modifications were made. As sequencing quality decreases for a 500-cycle kit, the sgRNA flanking sequences were extended to CTAGCTCTAAAAC and ACTAGTATTATAC. These flanking sequences were permitted to have up to 10 mismatches for sgRNAs 1 and 2, and 6 mismatches for sgRNA 3. Flanking sequences for the barcode were the same as above and were permitted to have up to 6 mismatches. The barcode itself was required to match a target exactly, however the cutoff to call a specific sgRNA homology region was lowered. Sequences in position 1 were allowed to have as many as 2 mismatches, and sequences in position 2 were allowed to have as many as 10 mismatches. This permissive filter was required due to the sharp decrease in sequencing confidence at the end of the sequencing run. However, if a sequenced sgRNA ambiguously mapped to multiple library sequences with the same number of mismatches, no identity was called. We performed next-generation sequencing analysis from HiSeq data on a high-performance computing cluster, while all subsequent analysis was performed locally.

We calculated relative growth rates following each CRISPRi treatment using the change in sgRNA counts over time. At each timepoint (t), we normalized counts for a specific barcoded CRISPRi construct (sgRNA) to the control construct (Nont) and our initial timepoint (t0) using Equation 4.

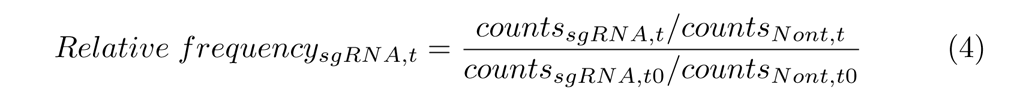

We then log_2_-transformed this relative frequency prior to growth rate fitting. For the pairwise sgRNA libraries, if fewer than 10 counts were identified for a construct at a given timepoint, no future timepoints were considered for that construct. Growth rates were calculated for CRISPRi constructs with sufficient counts over at least the first three timepoints. For pairwise and third-order CRISPRi experiments, the time axis was rescaled from hours to generations based on the overall culture generation time in each continuous culture vial. We then fit a line to our log_2_(Relative frequency) vs. generation data using scipy.stats.linregress and defined the slope of the best fit line as the raw growth rate. At this step, two sgRNAs (gdhA_1_42_B_MM14 and gdhA_3_216_B_MM8) were determined to have off-target effects and were removed from future analysis (see Jupyter Notebook 3_Pairwise_Growth_Rates.ipynb for details). For gene-by-environment analysis, continuous culture growth rates for each media and sgRNA combination were rescaled using plate reader absolute growth data using Equation 5.

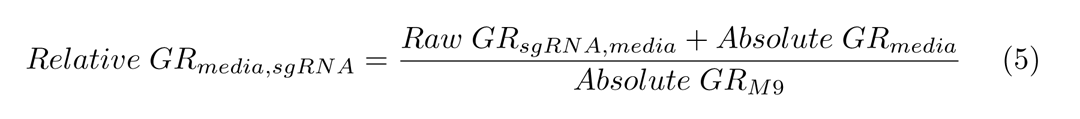

We then removed escapers, CRISPRi constructs that abnormally lacked a growth rate deficit, using a one-way Dixon Q-test for outliers at 95% confidence^33^. If at least four growth rates were successfully calculated across barcodes and sgRNA orders after removing escapers, the replicates were averaged to determine a mean relative growth rate. Finally, we normalized growth rates so the fully Nont control had a growth rate of 1 and the most severe growth rate perturbation observed (glyA_1_26_C + thyA_1_60_B_MM2, Min GR) had a growth rate of 0 using Equation 6.

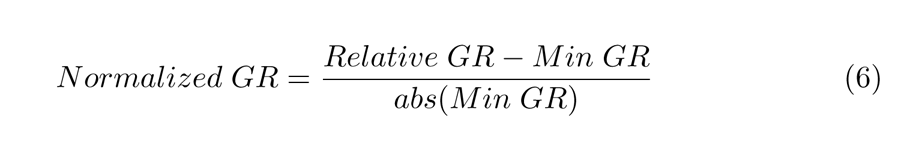

### Fitting the Coupling Model

We fit single-gene expression-growth rate curves using scipy.optimize.least_squares, bounding R_0_ to non-negative values with an initial guess of 0.5 for R_0_ and 0 for n. We fit gene-gene coupling values for each gene pair using the strategy described in Zimmer et al., 2016^28^, with some modifications. We optimized coupling constants a_ij_ and a_ji_ using scipy.optimize.least_squares, bounding between −1 and 10 with an initial guess of 0 for both. To limit the spread of coupling constant values, we calculated the RMSD between growth rate predictions and experimental measurements with an added regularization term λ * Σabs(a_ij_). We used a regularization value of λ = 10^-1.25^ after evaluating a range of values on subsampled data for both the pairwise and *folA*/*thyA* libraries (Fig S2). We calculated pairwise and third-order predicted growth rates using each gene’s individual repression-growth rate parameters and relevant coupling constants.

### Estimating Uncertainty with Bootstrapping

We used bootstrapping to computationally evaluate how sensitive each model’s predictions were to changes in single perturbation-growth rate fits. For each single perturbation curve, we generated a new training set of the same size as our input data set by resampling data with replacement from individual perturbation-growth rate measurements. For cases where both perturbation intensity and growth rate measurements had associated uncertainties (e.g., qPCR-based repression measurements and NGS-based growth rate measurements), we resampled these measurements independently. In all cases, we refit perturbation-growth curves using resampled data 100 times and used the resulting curves to estimate variance and establish 95% confidence intervals on our model’s fit parameters and accuracy (Table S2).

