## Supplemental Figures and Tables for "A continuous epistasis model for predicting growth rate given combinatorial variation in gene expression and environment"

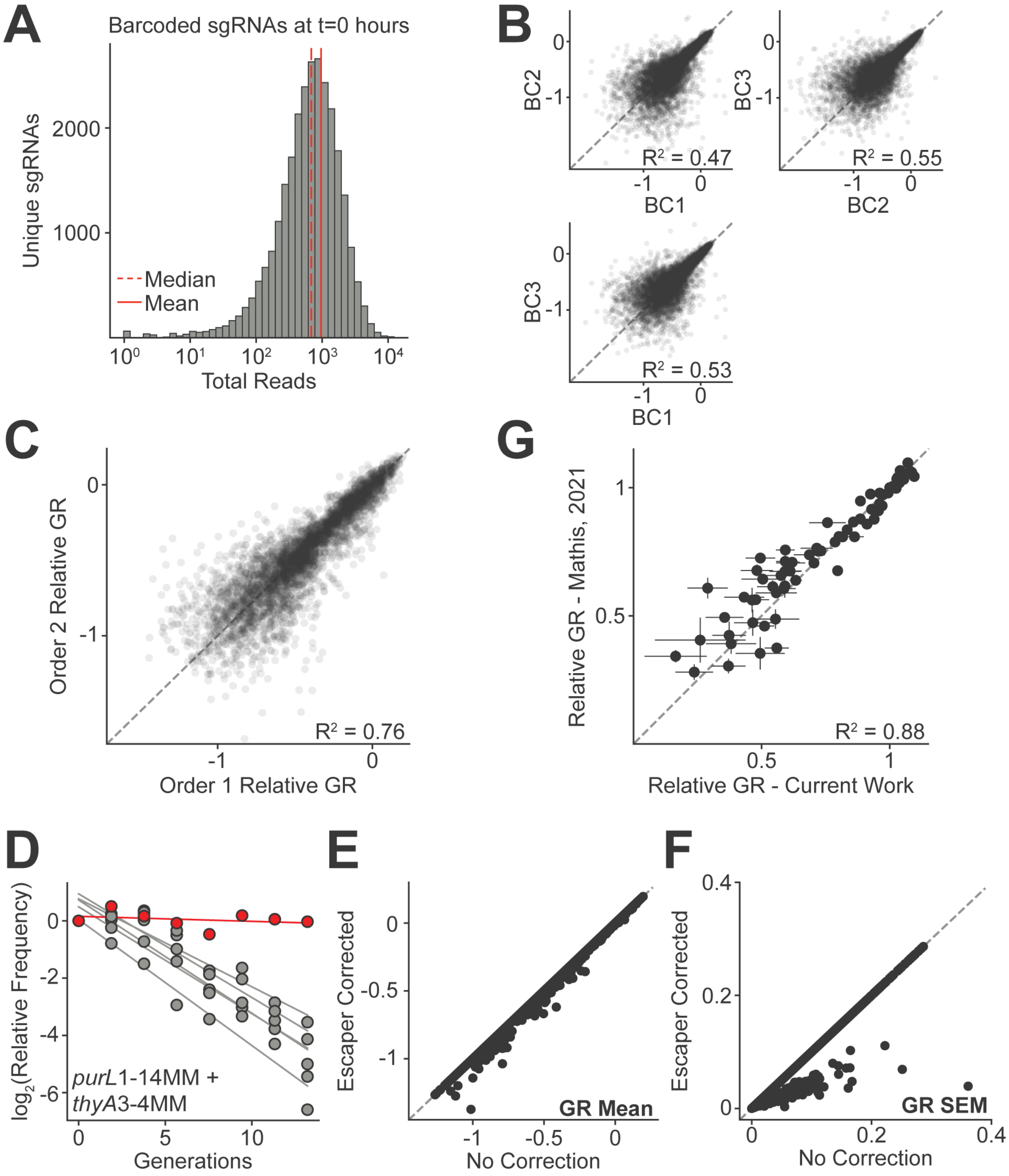

**Figure S1. Pairwise CRISPRi library completeness and quality control.** (A) Distribution of barcoded sgRNA sequencing counts at $T_0$ following filtering. The library has the expected log-normal distribution, with a mean of 964 counts (solid line) and a median of 683 counts (dotted line). (B) Growth rates from sgRNA combinations with a shared order (sgRNA1-sgRNA2) but different plasmid barcodes. As the data were normalized only following escaper correction and averaging, in these plots zero represents the growth rate of a non-targeting sgRNA control. (C) Growth rate correlation across sgRNA orders. Measurements are averaged across up to three barcoded replicates for each sgRNA order. (D) Six replicates of sgRNA relative frequency data for the *purL*1-14MM + *thyA*3-4MM sgRNA pair. These data contain an escaper, where one replicate (red) grows significantly faster than all other replicates (gray). (E) The effect of calculating growth rate with and without escaper correction. A growth rate of 0 represents the non-targeting sgRNA control. (F) The effect of escaper correction on the SEM of growth rate measurements. (G) Single CRISPRi perturbations from this study (one targeting and one non-targeting sgRNA) correlate well with those performed in an independent CRISPRi-growth rate experiment[S1] (one targeting sgRNA alone). Error bars represent the SEM of growth rate measurements across $n \geq 4$. The non-targeting sgRNA control's growth rate is normalized to 1, and 0 represents the slowest growth rate observed in the pairwise experiment. Dotted line is $y = x$ for all correlation plots.

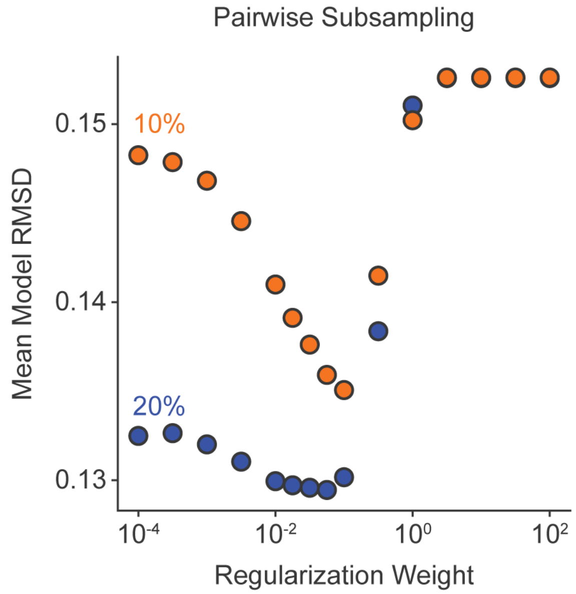

**Figure S2. Regularization optimization for pairwise expression predictions**. A range of regularization strengths were tested on subsampled data for the pairwise CRISPRi library. Either 10% (orange dots) or 20% (blue dots) of pairwise CRISPRi measurements were used to fit coupling constants, and error was quantified on the hold out data across 100 subsampling iterations. A final regularization weight of $10^{-1.25}$ was selected following this optimization.

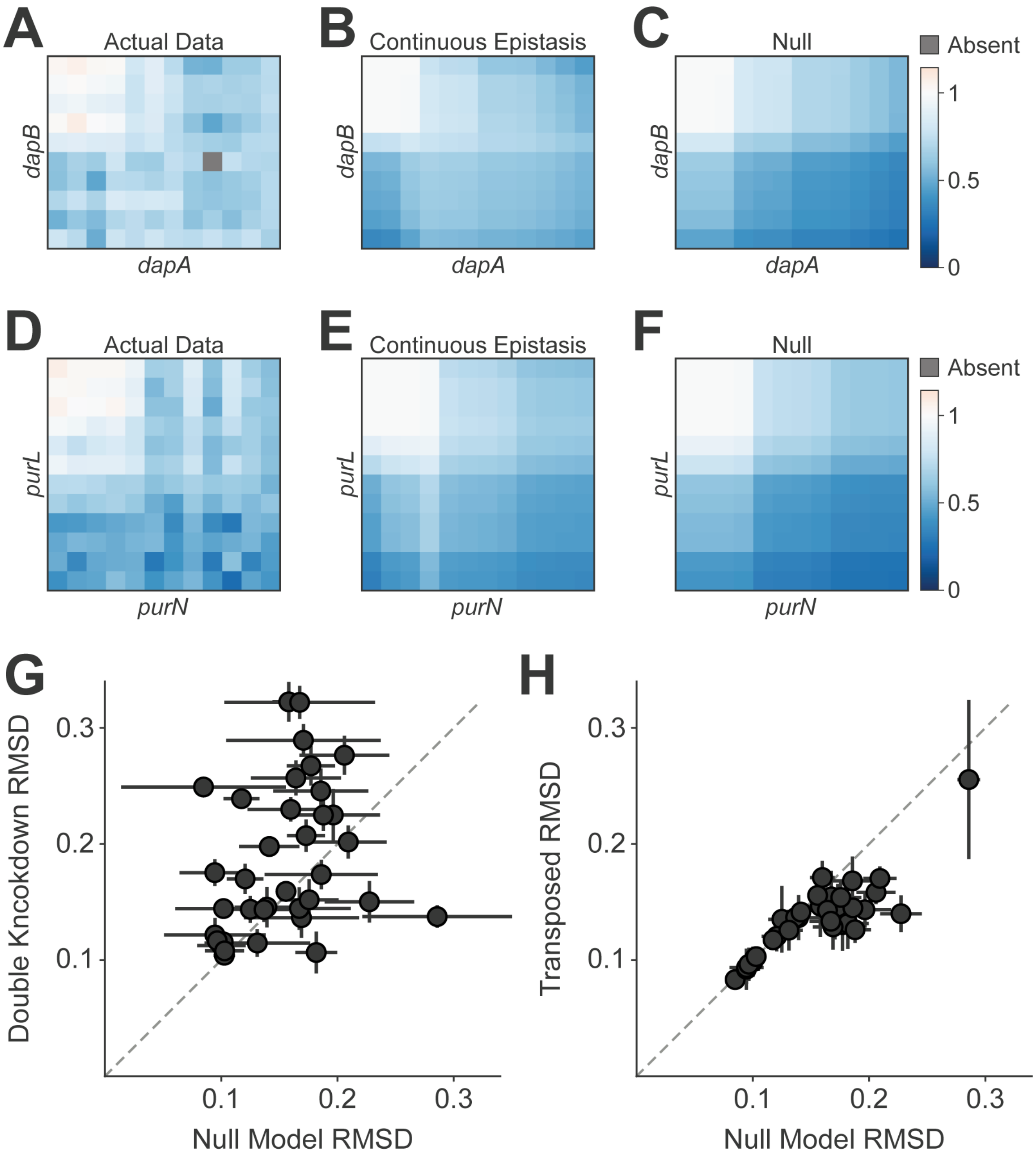

**Figure S3. The continuous epistasis model outperforms alternative models for predicting pairwise CRISPRi growth rates**. (A) Pairwise expression-growth rate data following CRISPRi knockdown of both *dapA* and *dapB*, which show positive coupling. Each row and column represents a unique sgRNA, and pixels represent the growth rate effect of a given sgRNA pair. Rows and columns are sorted by knockdown intensity, ranging from wildtype-like expression (top left) to maximal double-knockdown (bottom right). Gray indicates missing growth rate data. (B) Predicted expression-growth rate data utilizing the coupling-sensitive continuous epistasis model. (C) Predicted expression-growth rate data utilizing the coupling-insensitive Null model. (D-F) Same as A-C, but for the *purN*/*purL* gene pair, which shows no significant coupling. (G-H) Comparing the performance of the Null model to either (G) a model with couplings fit only on the most severe double-knockdown data or (H) a model with swapped $a_{ij}$ and $a_{ji}$ coupling constants, as in Fig 4G.

Error bars are standard deviations of models fit on bootstrapped single perturbation-growth curves. Dotted gray line is $y = x$.

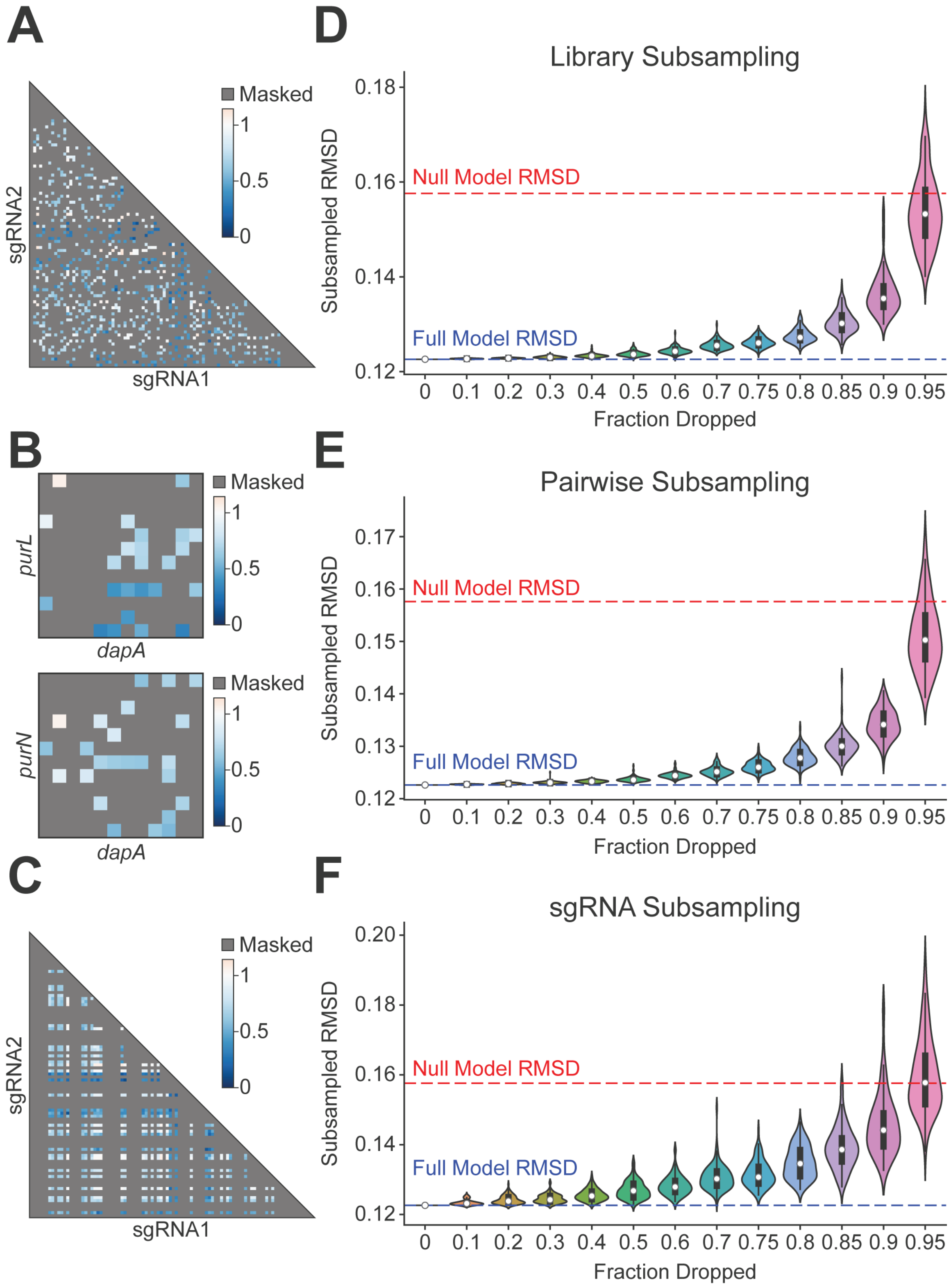

**Figure S4. Model performance is robust to subsampling the training data.** (A) A diagram of the library subsampling strategy, where 20% of pairwise sgRNA perturbations are randomly selected from the entire landscape. (B) A diagram of the pairwise subsampling strategy, where 20% of sgRNA perturbations are randomly selected from within each gene pair. (C) A diagram of the sgRNA subsampling strategy, where 20% of pairwise sgRNA perturbations are selected using a random subset of individual sgRNAs. (D-F) Each violin plot represents the average model RMSD across 100 subsampling iterations for its respective subsampling strategy after dropping the fraction of data indicated along the x-axis. Each white dot represents the median RMSD, and each wide gray bar represents the interquartile range. The dotted blue line represents the RMSD of the model fit on all pairwise data, while the dotted red line represents the RMSD of the Null model.

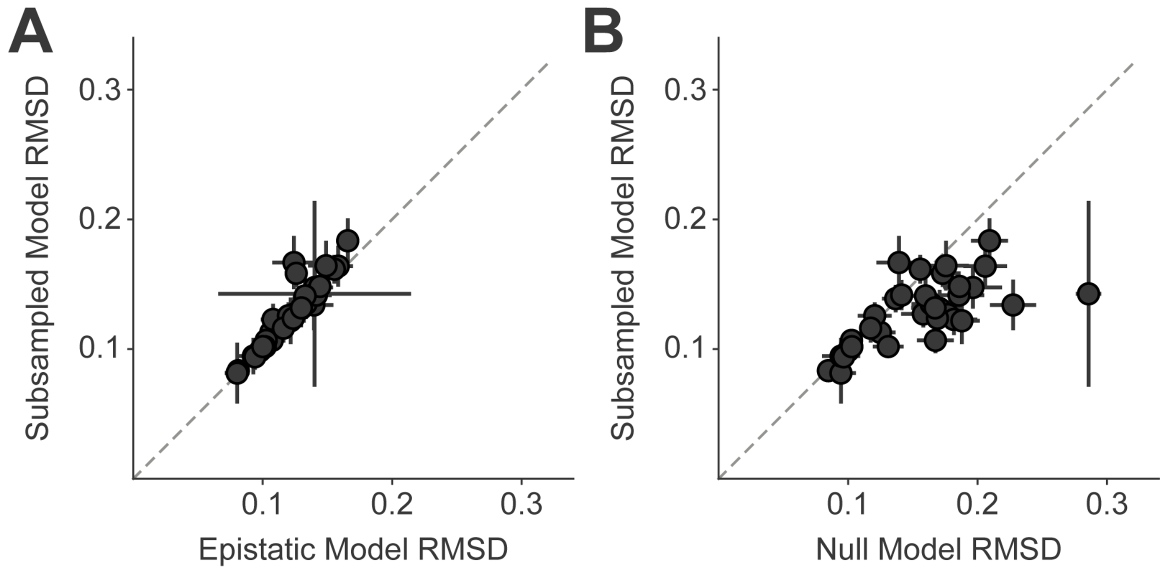

**Figure S5. Effect of subsampling pairwise growth rate data on the continuous epistasis model.** Subsampled model performance compared to the epistatic model without subsampling (A) or the Null model (B), as in Fig 4G. Error bars are standard deviations of models fit on bootstrapped single perturbation-growth curves. Dotted gray line is y = x.

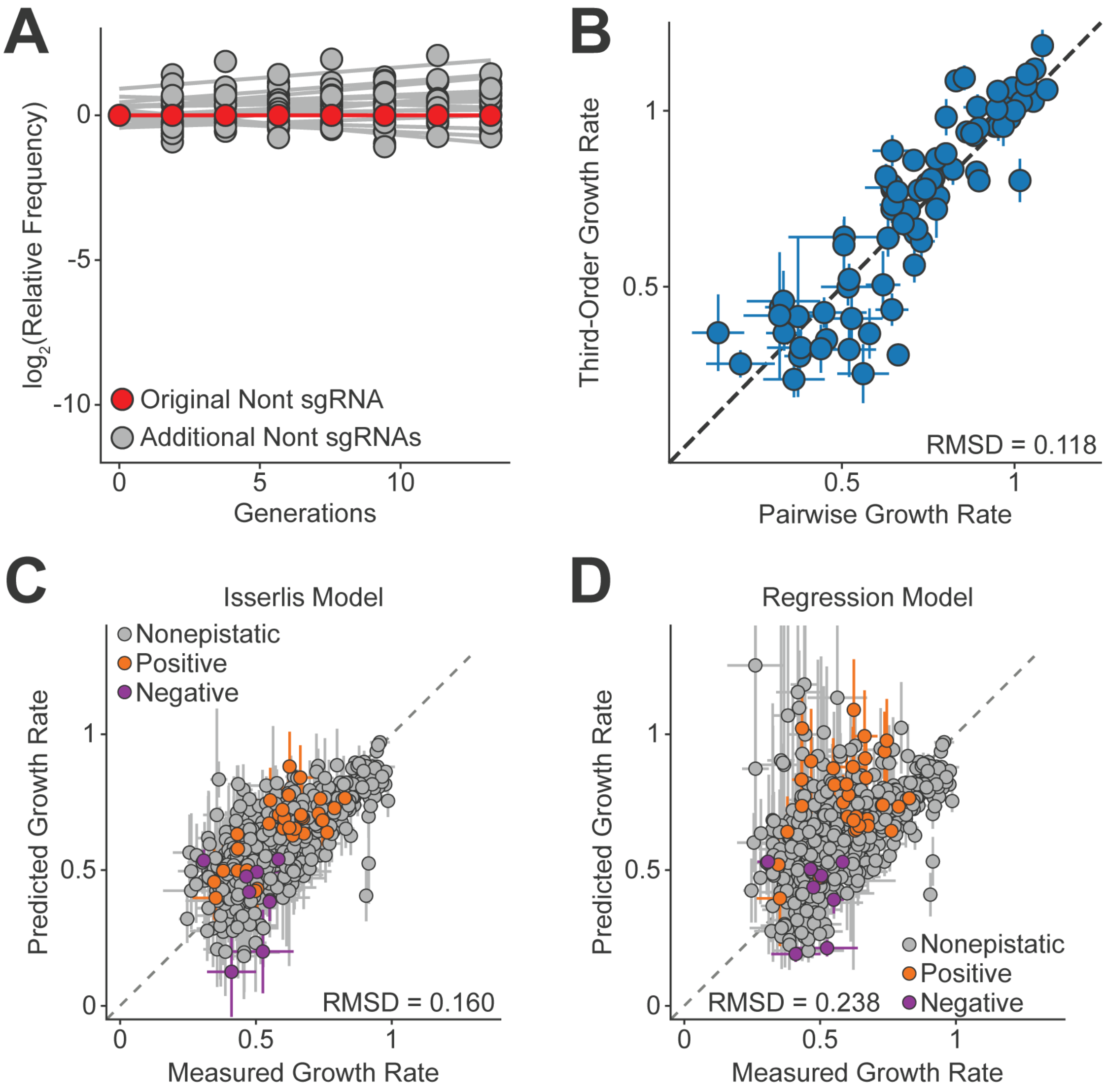

**Figure S6. The continuous epistasis model enhances predictions of higher-order combinations of gene knockdown.** (A) We validated multiple new non-targeting (Nont) controls (gray) relative to our previously validated control (red). The growth rate effect (slope of $\log_2$(relative frequency) over time) is near zero for all pairwise combinations of control sgRNAs. (B) Plotted are the growth rate effects of sgRNA constructs included in both the pairwise and third-order CRISPRi experiments. Error bars represent the SEM of growth rate measurements from $n \geq 4$ replicates. Dotted gray line is $y = x$. (C-D) Comparing model performance of the continuous epistasis model to the Isserlis (C) and regression (D) models, as in Fig 5C-D.

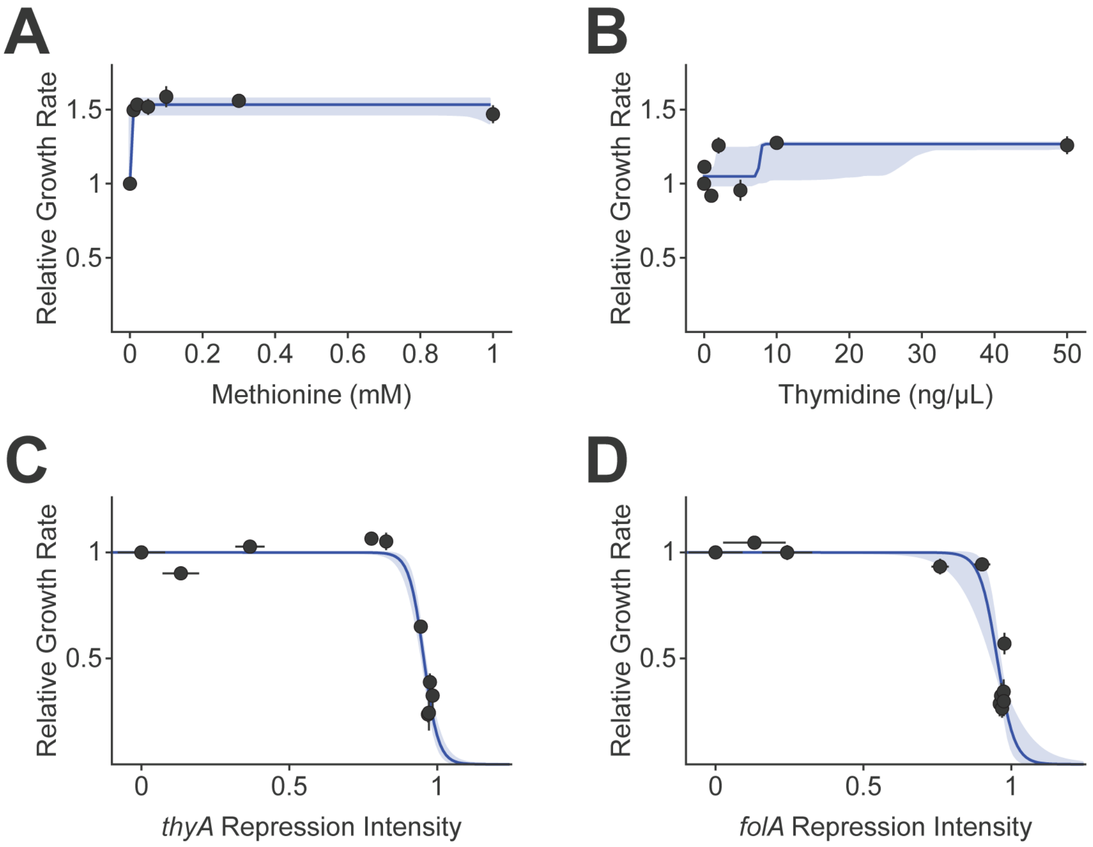

**Figure S7. Individual perturbation-growth rate relationships for repression and environment data.** (A-B) Supplement-growth rate relationships, where relative growth rates were measured by monitoring $OD_{600}$ over time in plate reader assays for methionine (A) and thymidine (B). Blue lines represent the four-parameter logistic fit to each data set, and the pale blue area represents a pointwise 95% confidence interval of this logistic fit. Vertical error bars represent SEM of growth rate measurements between n ≥ 4 replicates. (C-D) Expression-growth rate relationships for *thyA* (C) and *folA* (D), where relative growth rate measurements were made using an NGS-based growth rate assay. Error bars represent SEM of growth rate measurements between n ≥ 4 replicates (vertical) and RT-qPCR measurements across n ≥ 3 technical replicates (horizontal). Blue lines and the shaded area represent a two-parameter logistic fit and 95% confidence interval, respectively.

### Supplemental Tables

**Table S1. sgRNA insert and homology sequences.**

| sgRNA | Homology Sequence |
|---|---|
| dapA_1_98_C | AACGATCGCCGAAGTACCGC |
| dapA_1_98_W_MM1 | TACGATCGCCGAAGTACCGC |
| dapA_1_98_B_MM3 | TTGGATCGCCGAAGTACCGC |
| dapA_1_98_B_MM7 | TTGCTAGGCCGAAGTACCGC |
| dapA_1_98_B_MM9 | TTGCTAGCGCGAAGTACCGC |
| dapA_1_98_B_MM14 | TTGCTAGCGGCTTCTACCGC |
| dapA_3_214_C | GCGCCGGTCCCGGCAATTAC |
| dapA_3_214_W_MM1 | CCGCCGGTCCCGGCAATTAC |
| dapA_3_214_B_MM6 | CGCGGCGTCCCGGCAATTAC |
| dapA_3_214_B_MM9 | CGCGGCCAGCCGGCAATTAC |
| dapA_3_214_B_MM14 | CGCGGCCAGGGCCGAATTAC |
| dapB_1_18_C | CGGCTCCCGCGATGGCAACG |
| dapB_1_18_B_MM5 | GCCGACCCGCGATGGCAACG |
| dapB_1_18_B_MM11 | GCCGAGGGCGCATGGCAACG |
| dapB_1_18_B_MM13 | GCCGAGGGCGCTAGGCAACG |
| dapB_1_18_B_MM14 | GCCGAGGGCGCTACGCAACG |
| dapB_3_239_C | GTTCAGCGTACCTTCCGGAC |
| dapB_3_239_W_MM1 | CTTCAGCGTACCTTCCGGAC |
| dapB_3_239_B_MM9 | CAAGTCGCAACCTTCCGGAC |
| dapB_3_239_B_MM14 | CAAGTCGCATGGAACCGGAC |
| gdhA_1_42_C | TTTGATTCGGGTCGCGCTTT |
| gdhA_1_42_B_MM4 | AAACATTCGGGTCGCGCTTT |
| gdhA_1_42_B_MM6 | AAACTATCGGGTCGCGCTTT |
| gdhA_1_42_B_MM8 | AAACTAAGGGGTCGCGCTTT |
| gdhA_1_42_B_MM10 | AAACTAAGCCGTCGCGCTTT |
| gdhA_1_42_B_MM14 | AAACTAAGCCCAGCCGCTTT |
| gdhA_3_216_C | ATGCACGGTTGACCTGTATC |
| gdhA_3_216_B_MM4 | TACGACGGTTGACCTGTATC |
| gdhA_3_216_B_MM6 | TACGTGGGTTGACCTGTATC |
| gdhA_3_216_B_MM8 | TACGTGCCTTGACCTGTATC |
| gdhA_3_216_B_MM10 | TACGTGCCAAGACCTGTATC |
| gdhA_3_216_B_MM14 | TACGTGCCAACTGGTGTATC |
| gltB_1_17_C | ACCACAGTTATCCCTCTCAA |
| gltB_1_17_B_MM4 | TGGTCAGTTATCCCTCTCAA |
| gltB_1_17_B_MM6 | TGGTGTGTTATCCCTCTCAA |
| gltB_1_17_B_MM8 | TGGTGTCATATCCCTCTCAA |
| gltB_1_17_B_MM10 | TGGTGTCAATTCCCTCTCAA |
| gltB_1_17_B_MM14 | TGGTGTCAATAGGGTCTCAA |
| gltB_3_284_C | AACGATGCGGCGTGCGGCAG |
| gltB_3_284_B_MM4 | TTGCATGCGGCGTGCGGCAG |
| gltB_3_284_B_MM6 | TTGCTAGCGGCGTGCGGCAG |
| gltB_3_284_B_MM8 | TTGCTACGGGCGTGCGGCAG |
| gltB_3_284_B_MM10 | TTGCTACGCCCGTGCGGCAG |
| gltB_3_284_B_MM14 | TTGCTACGCCGCACCGGCAG |
| folA_1_56_C | GGCAGGCAGGTTCCACGGCA |
| folA_1_56_B_MM2 | CCCAGGCAGGTTCCACGGCA |
| folA_1_56_B_MM3 | CCGAGGCAGGTTCCACGGCA |

| | |
|---|---|
| folA_1_56_B_MM6 | CCGTCCCAGGTTCCACGGCA |
| folA_1_56_B_MM9 | CCGTCCGTCGTTCCACGGCA |
| folA_1_56_B_MM10 | CCGTCCGTCCTTCCACGGCA |
| folA_1_56_B_MM11 | CCGTCCGTCCATCCACGGCA |
| folA_1_56_B_MM14 | CCGTCCGTCCAAGGACGGCA |
| folA_3_242_C | TACGTCACCACACGCCGCGA |
| folA_3_242_B_MM3 | ATGGTCACCACACGCCGCGA |
| thyA_1_60_C | AAAGCGTTCCGGTTCCGGTA |
| thyA_1_60_B_MM2 | TTAGCGTTCCGGTTCCGGTA |
| thyA_1_60_B_MM12 | TTTCGCAAGGCCTTCCGGTA |
| thyA_1_60_B_MM13 | TTTCGCAAGGCCATCCGGTA |
| thyA_3_233_C | ATCGGCCCATTCGTCCCAGA |
| thyA_3_233_B_MM4 | TAGCGCCCATTCGTCCCAGA |
| thyA_3_233_B_MM5 | TAGCCCCCATTCGTCCCAGA |
| thyA_3_233_B_MM6 | TAGCCGCCATTCGTCCCAGA |
| thyA_3_233_B_MM7 | TAGCCGGCATTCGTCCCAGA |
| glyA_1_26_C | CCACAGTTCGGCATCATAAT |
| glyA_1_26_B_MM4 | GGTGAGTTCGGCATCATAAT |
| glyA_1_26_B_MM8 | GGTGTCAACGGCATCATAAT |
| glyA_1_26_B_MM9 | GGTGTCAAGGGCATCATAAT |
| glyA_1_26_B_MM11 | GGTGTCAAGCCCATCATAAT |
| glyA_1_26_B_MM12 | GGTGTCAAGCCGATCATAAT |
| glyA_3_279_C | CCTGGGAGCCGGAGTGCGGC |
| glyA_3_279_B_MM3 | GGAGGGAGCCGGAGTGCGGC |
| glyA_3_279_B_MM6 | GGACCCAGCCGGAGTGCGGC |
| glyA_3_279_B_MM14 | GGACCCTCGGCCTCTGCGGC |
| purN_1_86_C | ATTGCTGAAAACTGCCCGTA |
| purN_1_86_B_MM2 | TATGCTGAAAACTGCCCGTA |
| purN_1_86_B_MM4 | TAACCTGAAAACTGCCCGTA |
| purN_1_86_B_MM12 | TAACGACTTTTGTGCCCGTA |
| purN_3_238_C | CCAGCCAGCACGACCACATC |
| purN_3_238_W_MM1 | GCAGCCAGCACGACCACATC |
| purN_3_238_B_MM4 | GGTCCCAGCACGACCACATC |
| purN_3_238_B_MM5 | GGTCGCAGCACGACCACATC |
| purN_3_238_B_MM10 | GGTCGGTCGTCGACCACATC |
| purN_3_238_B_MM11 | GGTCGGTCGTGGACCACATC |
| purN_3_238_B_MM14 | GGTCGGTCGTGCTGCACATC |
| purL_1_25_C | ATTCGGAATGCCGACAGTGC |
| purL_1_25_B_MM3 | TAACGGAATGCCGACAGTGC |
| purL_1_25_B_MM8 | TAAGCCTTTGCCGACAGTGC |
| purL_1_25_B_MM9 | TAAGCCTTAGCCGACAGTGC |
| purL_1_25_B_MM14 | TAAGCCTTACGGCTCAGTGC |
| purL_3_201_C | GGAGTTTGCCTTGCGGGGCG |
| purL_3_201_W_MM1 | CGAGTTTGCCTTGCGGGGCG |
| purL_3_201_B_MM5 | CCTCATTGCCTTGCGGGGCG |
| purL_3_201_B_MM9 | CCTCAAACGCTTGCGGGGCG |
| purL_3_201_B_MM11 | CCTCAAACGGATGCGGGGCG |
| purL_3_201_B_MM14 | CCTCAAACGGAACGGGGGCG |
| non-targeting | AGTAATTGTATAGGCGACCT |

The full sgRNA insert sequence is provided below. The string of “X”s in the sequence indicate location of the sgRNA homology sequence.

TTGACAGCTAGCTCAGTCCTAGGTATAATACTAGTXXXXXXXXXXXXXXXXXXXXGTTTTAGAGCTAGAAA
TAGCAAGTTAAAATAAGGCTAGTCCGTTATCAACTTGAAAAAGTGGCACCGAGTCGGTGCTTTTTTTGAAG

**Table S2. Single CRISPRi knockdown expression-growth rate curves.**

| Gene | $R_0$ | $R_0$ 95% CI | Steepness | Steepness 95% CI |
|---|---|---|---|---|
| *dapA* | 0.852 | [0.817, 0.934] | 6.78 | [4.057, 8.209] |
| *dapB* | 0.91 | [0.863, 1.069] | 5.511 | [2.418, 8.216] |
| *gdhA* | 1.142 | [1.056, 3.185] | 20.536 | [2.729, 52.207] |
| *gltB* | 2.262 | [1.289, 7.248] | 1.704 | [0.353, 7.313] |
| *folA* | 0.951 | [0.93, 0.961] | 36.048 | [18.245, 70.715] |
| *thyA* | 0.972 | [0.964, 0.978] | 37.296 | [30.659, 46.881] |
| *glyA* | 0.989 | [0.977, 1.192] | 27.761 | [3.381, 44.626] |
| *purN* | 0.998 | [0.984, 1.016] | 21.852 | [16.244, 34.591] |
| *purL* | 0.975 | [0.968, 0.995] | 23.274 | [10.72, 37.373] |

**Table S3. Epistatic couplings between pairwise CRISPRi knockdowns.**

| Gene 1 | Gene 2 | $a_{ij}$ | $a_{ij}$ 95% CI | $a_{ji}$ | $a_{ji}$ 95% CI | Model RMSD | Model RMSD 95% CI | Null RMSD | Null RMSD 95% CI |
|---|---|---|---|---|---|---|---|---|---|
| *dapA* | *dapB* | 0.361 | [0.237, 0.449] | 0.327 | [0.267, 0.387] | 0.14 | [0.123, 0.178] | 0.227 | [0.192, 0.26] |
| *dapA* | *gdhA* | 0 | [-0.0, 0.014] | 0.049 | [0.0, 0.093] | 0.081 | [0.076, 0.093] | 0.085 | [0.076, 0.105] |
| *dapA* | *gltB* | 0 | [-0.0, 0.08] | 0.108 | [0.0, 0.211] | 0.08 | [0.068, 0.105] | 0.094 | [0.075, 0.119] |
| *dapA* | *folA* | 0.04 | [0.024, 0.079] | 0.082 | [-0.0, 0.221] | 0.124 | [0.119, 0.15] | 0.158 | [0.138, 0.194] |
| *dapA* | *thyA* | 0.022 | [0.015, 0.037] | 0.072 | [0.0, 0.162] | 0.106 | [0.102, 0.124] | 0.125 | [0.107, 0.151] |
| *dapA* | *glyA* | 0.071 | [0.049, 0.261] | 0.095 | [0.036, 0.183] | 0.14 | [0.117, 0.169] | 0.196 | [0.155, 0.236] |
| *dapA* | *purN* | 0.061 | [0.042, 0.099] | 0.197 | [0.132, 0.238] | 0.107 | [0.096, 0.123] | 0.167 | [0.138, 0.19] |
| *dapA* | *purL* | 0.069 | [0.046, 0.135] | 0.059 | [0.033, 0.076] | 0.13 | [0.113, 0.154] | 0.171 | [0.149, 0.205] |
| *dapB* | *gdhA* | 0 | [-0.0, 0.027] | 0.024 | [0.0, 0.078] | 0.093 | [0.07, 0.121] | 0.095 | [0.07, 0.127] |
| *dapB* | *gltB* | 0 | [-0.0, 0.128] | 0.132 | [0.0, 0.265] | 0.124 | [0.103, 0.164] | 0.139 | [0.107, 0.176] |
| *dapB* | *folA* | 0.066 | [0.044, 0.133] | 0.095 | [0.0, 0.231] | 0.124 | [0.117, 0.154] | 0.177 | [0.151, 0.216] |
| *dapB* | *thyA* | 0.057 | [0.045, 0.085] | 0.127 | [0.0, 0.268] | 0.126 | [0.12, 0.147] | 0.173 | [0.151, 0.204] |
| *dapB* | *glyA* | 0.088 | [0.06, 0.262] | 0.088 | [0.0, 0.185] | 0.158 | [0.14, 0.181] | 0.206 | [0.172, 0.237] |
| *dapB* | *purN* | 0.1 | [0.066, 0.161] | 0.222 | [0.078, 0.303] | 0.108 | [0.096, 0.128] | 0.182 | [0.144, 0.211] |
| *dapB* | *purL* | 0.123 | [0.088, 0.235] | 0.084 | [0.0, 0.123] | 0.122 | [0.105, 0.147] | 0.188 | [0.161, 0.216] |
| *gdhA* | *gltB* | 0 | [-1.063, 0.0] | -0.425 | [-1.19, 0.0] | 0.14 | [0.088, 0.31] | 0.286 | [0.273, 0.312] |
| *gdhA* | *folA* | 0.004 | [-0.005, 0.044] | 0 | [-0.0, 0.004] | 0.12 | [0.115, 0.151] | 0.121 | [0.115, 0.159] |
| *gdhA* | *thyA* | 0.006 | [-0.003, 0.02] | 0.002 | [-0.0, 0.042] | 0.098 | [0.093, 0.116] | 0.1 | [0.094, 0.126] |
| *gdhA* | *glyA* | 0.003 | [-0.017, 0.02] | 0 | [-0.0, 0.0] | 0.155 | [0.155, 0.174] | 0.156 | [0.156, 0.181] |
| *gdhA* | *purN* | 0 | [-0.017, 0.009] | 0 | [-0.0, 0.0] | 0.102 | [0.102, 0.111] | 0.102 | [0.102, 0.114] |
| *gdhA* | *purL* | 0 | [-0.01, 0.012] | 0 | [-0.0, 0.0] | 0.102 | [0.099, 0.126] | 0.102 | [0.099, 0.126] |
| *gltB* | *folA* | 0.008 | [-0.004, 0.045] | 0 | [-0.0, 0.0] | 0.116 | [0.114, 0.139] | 0.117 | [0.117, 0.141] |
| *gltB* | *thyA* | 0 | [-0.017, 0.012] | 0 | [-0.0, 0.0] | 0.103 | [0.102, 0.114] | 0.103 | [0.102, 0.118] |
| *gltB* | *glyA* | 0.017 | [0.0, 0.055] | 0 | [-0.0, 0.0] | 0.132 | [0.127, 0.156] | 0.137 | [0.128, 0.164] |
| *gltB* | *purN* | 0.013 | [0.0, 0.048] | 0 | [-0.0, 0.004] | 0.094 | [0.092, 0.108] | 0.097 | [0.093, 0.111] |

| | | | | | | | | | |
|---|---|---|---|---|---|---|---|---|---|
| *gltB* | *purL* | 0.012 | [0.0, 0.022] | 0 | [-0.0, 0.0] | 0.102 | [0.1, 0.116] | 0.103 | [0.102, 0.117] |
| *folA* | *thyA* | 0.028 | [0.012, 0.055] | 0.047 | [0.025, 0.104] | 0.166 | [0.164, 0.178] | 0.209 | [0.191, 0.242] |
| *folA* | *glyA* | 0.073 | [0.0, 0.16] | 0.023 | [0.012, 0.059] | 0.141 | [0.132, 0.174] | 0.186 | [0.154, 0.231] |
| *folA* | *purN* | 0.026 | [-0.0, 0.066] | 0.045 | [0.027, 0.076] | 0.124 | [0.12, 0.143] | 0.169 | [0.148, 0.208] |
| *folA* | *purL* | 0.052 | [0.019, 0.101] | 0.026 | [0.016, 0.044] | 0.133 | [0.122, 0.159] | 0.164 | [0.149, 0.203] |
| *thyA* | *glyA* | 0 | [-0.035, 0.022] | 0 | [-0.007, 0.0] | 0.141 | [0.137, 0.159] | 0.141 | [0.14, 0.161] |
| *thyA* | *purN* | 0.045 | [0.016, 0.071] | 0.036 | [0.023, 0.053] | 0.144 | [0.14, 0.155] | 0.186 | [0.166, 0.215] |
| *thyA* | *purL* | 0.058 | [0.037, 0.12] | 0.009 | [0.004, 0.018] | 0.133 | [0.126, 0.153] | 0.16 | [0.148, 0.187] |
| *glyA* | *purN* | 0.027 | [0.017, 0.063] | 0.046 | [0.012, 0.132] | 0.13 | [0.119, 0.148] | 0.167 | [0.135, 0.2] |
| *glyA* | *purL* | 0.04 | [0.028, 0.079] | 0.022 | [0.011, 0.053] | 0.149 | [0.128, 0.179] | 0.176 | [0.145, 0.211] |
| *purN* | *purL* | 0.053 | [0.034, 0.095] | 0.009 | [0.002, 0.016] | 0.1 | [0.094, 0.117] | 0.131 | [0.113, 0.158] |

**Table S4. Single genetic and environmental perturbation curves.**

| Perturbation | $D_0$ | $D_0$ 95% CI | Steepness | Steepness 95% CI | $g_{max}$ | $g_{max}$ 95% CI | $g_{min}$ | $g_{min}$ 95% CI |
|---|---|---|---|---|---|---|---|---|
| thymidine | 7.63 | [1.175, 28.655] | -8.373 | [-13.991, -0.166] | 1.267 | [1.239, 1.275] | 1.049 | [0.919, 1.093] |
| methionine | 0.008 | [0.004, 16.543] | -1562.1 | [-2749.892, 553.582] | 1.533 | [1.008, 1.587] | 1 | [1.0, 1.541] |
| folA_KD | 0.952 | [0.934, 0.961] | 34.401 | [15.616, 56.3] | N/A | N/A | N/A | N/A |
| thyA_KD | 0.955 | [0.949, 0.958] | 40.92 | [32.275, 51.489] | N/A | N/A | N/A | N/A |

**Table S5. Epistatic couplings between CRISPRi knockdowns and environmental shifts.**

| Perturbation 1 | Perturbation 2 | Perturbation 3 | Perturbation 4 | $a_{ij}$ | $a_{ij}$ 95% CI | $a_{ji}$ | $a_{ji}$ 95% CI | Model RMSD | Model RMSD 95% CI | Null RMSD | Null RMSD 95% CI |
|---|---|---|---|---|---|---|---|---|---|---|---|
| thymidine | methionine | N/A | N/A | 0 | [-0.0, 0.0] | 0 | [0.0, 0.141] | 0.151 | [0.138, 0.227] | 0.151 | [0.138, 0.251] |
| thymidine | folA_KD | N/A | N/A | 0 | [-0.0, 0.0] | 0.047 | [0.017, 0.127] | 0.17 | [0.168, 0.2] | 0.235 | [0.209, 0.272] |
| thymidine | thyA_KD | N/A | N/A | 0 | [-0.0, 0.0] | 0.73 | [0.197, 1.288] | 0.118 | [0.115, 0.24] | 0.536 | [0.508, 0.573] |
| methionine | folA_KD | N/A | N/A | 0 | [-0.0, 0.0] | 0.013 | [0.0, 0.047] | 0.151 | [0.146, 0.208] | 0.184 | [0.157, 0.239] |
| methionine | thyA_KD | N/A | N/A | 0 | [-0.0, 0.0] | 0.009 | [0.0, 0.027] | 0.167 | [0.16, 0.195] | 0.183 | [0.166, 0.216] |
| folA_KD | thyA_KD | N/A | N/A | 0.011 | [-0.003, 0.037] | 0.009 | [0.0, 0.022] | 0.121 | [0.12, 0.133] | 0.125 | [0.122, 0.141] |
| thymidine | methionine | folA_KD | N/A | N/A | N/A | N/A | N/A | 0.214 | [0.206, 0.276] | 0.286 | [0.247, 0.344] |
| thymidine | methionine | thyA_KD | N/A | N/A | N/A | N/A | N/A | 0.194 | [0.189, 0.334] | 0.642 | [0.596, 0.697] |
| thymidine | folA_KD | thyA_KD | N/A | N/A | N/A | N/A | N/A | 0.262 | [0.243, 0.336] | 0.606 | [0.583, 0.63] |
| methionine | folA_KD | thyA_KD | N/A | N/A | N/A | N/A | N/A | 0.192 | [0.188, 0.238] | 0.256 | [0.235, 0.287] |
| thymidine | methionine | folA_KD | thyA_KD | N/A | N/A | N/A | N/A | 0.337 | [0.294, 0.451] | 0.727 | [0.689, 0.761] |

**Table S6. Primer sequences.**

| Name | Sequence | Reaction | Comments |
|---|---|---|---|
| GG_UnivF | GTACTGGGTCTCTAGGTTTGACAGCTAGCTCAGTCCTAG | Golden Gate Assembly | |
| GG_R1 | GTACTGGGTCTCTGGTACTTCAAAAAAAGCACCGACTCG | Golden Gate Assembly | |
| GG_F2 | GTACTGGGTCTCTTACCTTGACAGCTAGCTCAGTCCTAG | Golden Gate Assembly | |
| GG_R2 | GTACTGGGTCTCTACAACTTCAAAAAAAGCACCGACTCG | Golden Gate Assembly | |
| GG_F3 | GTACTGGGTCTCTTTGTTTGACAGCTAGCTCAGTCCTAG | Golden Gate Assembly | |
| GG_UnivR | GTACTGGGTCTCTCCACCTTCAAAAAAAGCACCGACTCG | Golden Gate Assembly | |
| TruSeqF | TGACTGGAGTTCAGACGTGTGCTCTTCCGATCTNNNNCTTCACCTCGAGAGGTTTGACAG | TruSeq amplification | N = Random nucleotide |
| TruSeqR_trunc | CACTCTTTCCCTACACGACGCTCTTCCGATCTNNNNTATTAGTACAGCGAGGCAAC | TruSeq amplification | N = Random nucleotide |
| TruSeqR | CACTCTTTCCCTACACGACGCTCTTCCGATCTNNNNCCATTATTAGTACAGCGAGGCAAC | TruSeq amplification | N = Random nucleotide |
| i5 Primer | AATGATACGGCGACCACCGAGATCTACACXXXXXXXXACACTCTTTCCCTACACGACGCTCTTCCGATCT | Illumina Index addition | X = Barcode nucleotide |
| i7 Primer | CAAGCAGAAGACGGCATACGAGATXXXXXXXXGTGACTGGAGTTCAGACGTGTGCTCTTCCGATC | Illumina Index addition | X = Barcode nucleotide |
| CPCR_pCRISPR_SeqF2 | GAACAGGAGAGCGCACGAGG | Colony PCR | |
| CPCR_pCRISPR_SeqR | CCATCAGATCCTTGGCGGC | Colony PCR | |
| iPCR_UnivR | ACTAGTATTATACCTAGGACTGAGCTAGC | iPCR | |
| iPCR_dapA_1_98_C_F | AACGATCGCCGAAGTACCGCGTTTTAGAGCTAGAAATAGCAAGTTAAAATAAGGC | iPCR | |
| iPCR_dapA_3_214_W_MM1_F | CCGCCGGTCCCGGCAATTACGTTTTAGAGCTAGAAATAGCAAGTTAAAATAAGGC | iPCR | |
| iPCR_dapA_3_214_B_MM9_F | CGCGGCCAGCCGGCAATTACGTTTTAGAGCTAGAAATAGCAAGTTAAAATAAGGC | iPCR | |
| iPCR_dapB_1_18_C_F | CGGCTCCCGCGATGGCAACGGTTTTAGAGCTAGAAATAGCAAGTTAAAATAAGGC | iPCR | |
| iPCR_dapB_1_18_B_MM11_F | GCCGAGGGCGCATGGCAACGGTTTTAGAGCTAGAAATAGCAAGTTAAAATAAGGC | iPCR | |
| iPCR_dapB_3_239_B_MM9_F | CAAGTCGCAACCTTCCGGACGTTTTAGAGCTAGAAATAGCAAGTTAAAATAAGGC | iPCR | |
| iPCR_gdhA_1_42_C_F | TTTGATTCGGGTCGCGCTTTGTTTTAGAGCTAGAAATAGCAAGTTAAAATAAGGC | iPCR | |
| iPCR_gdhA_1_42_B_MM6 | AAACTATCGGGTCGCGCTTTGTTTTAGAGCTAGAAATAGCAAGTTAAAATAAGGC | iPCR | |
| iPCR_gdhA_1_42_B_MM8_F | AAACTAAGGGGTCGCGCTTTGTTTTAGAGCTAGAAATAGCAAGTTAAAATAAGGC | iPCR | |
| iPCR_gdhA_1_42_B_MM14_F | AAACTAAGCCCAGCCGCTTTGTTTTAGAGCTAGAAATAGCAAGTTAAAATAAGGC | iPCR | |
| iPCR_gdhA_3_216_B_MM4_F | TACGACGGTTGACCTGTATCGTTTTAGAGCTAGAAATAGCAAGTTAAAATAAGGC | iPCR | |
| iPCR_gltB_1_17_C_F | ACCACAGTTATCCCTCTCAAGTTTTAGAGCTAGAAATAGCAAGTTAAAATAAGGC | iPCR | |

| | | |
|---|---|---|
| iPCR_gltB_1_17_B_MM6_F | TGGTGTGTTATCCCTCTCAAGTTTTAGAGCTAGAAATAGCAAGTTAAAATAAGGC | iPCR |
| iPCR_gltB_1_17_B_MM8_F | TGGTGTCATATCCCTCTCAAGTTTTAGAGCTAGAAATAGCAAGTTAAAATAAGGC | iPCR |
| iPCR_gltB_3_284_C_F | AACGATGCGGCGTGCGGCAGGTTTTAGAGCTAGAAATAGCAAGTTAAAATAAGGC | iPCR |
| iPCR_gltB_3_284_B_MM8_F | TTGCTACGGGCGTGCGGCAGGTTTTAGAGCTAGAAATAGCAAGTTAAAATAAGGC | iPCR |
| iPCR_gltB_3_284_B_MM14_F | TTGCTACGCCGCACCGGCAGGTTTTAGAGCTAGAAATAGCAAGTTAAAATAAGGC | iPCR |
| iPCR_folA_1_56_B_MM3_F | CCGAGGCAGGTTCCACGGCAGTTTTAGAGCTAGAAATAGCAAGTTAAAATAAGGC | iPCR |
| iPCR_folA_1_56_B_MM10_F | CCGTCCGTCCTTCCACGGCAGTTTTAGAGCTAGAAATAGCAAGTTAAAATAAGGC | iPCR |
| iPCR_thyA_1_60_B_MM13_F | TTTCGCAAGGCCATCCGGTAGTTTTAGAGCTAGAAATAGCAAGTTAAAATAAGGC | iPCR |
| iPCR_thyA_3_233_C_F | ATCGGCCCATTCGTCCCAGAGTTTTAGAGCTAGAAATAGCAAGTTAAAATAAGGC | iPCR |
| iPCR_thyA_3_233_B_MM5_F | TAGCCCCCATTCGTCCCAGAGTTTTAGAGCTAGAAATAGCAAGTTAAAATAAGGC | iPCR |
| iPCR_purN_1_86_C_F | ATTGCTGAAAACTGCCCGTAGTTTTAGAGCTAGAAATAGCAAGTTAAAATAAGGC | iPCR |
| iPCR_purN_1_86_B_MM2_F | TATGCTGAAAACTGCCCGTAGTTTTAGAGCTAGAAATAGCAAGTTAAAATAAGGC | iPCR |
| iPCR_purN_1_86_B_MM4_F | TAACCTGAAAACTGCCCGTAGTTTTAGAGCTAGAAATAGCAAGTTAAAATAAGGC | iPCR |
| iPCR_purN_1_86_B_MM12_F | TAACGACTTTTGTGCCCGTAGTTTTAGAGCTAGAAATAGCAAGTTAAAATAAGGC | iPCR |
| iPCR_purN_3_238_B_MM11_F | GGTCGGTCGTGGACCACATCGTTTTAGAGCTAGAAATAGCAAGTTAAAATAAGGC | iPCR |
| iPCR_purN_3_238_B_MM14_F | GGTCGGTCGTGCTGCACATCGTTTTAGAGCTAGAAATAGCAAGTTAAAATAAGGC | iPCR |
| iPCR_purL_1_25_B_MM9_F | TAAGCCTTAGCCGACAGTGCGTTTTAGAGCTAGAAATAGCAAGTTAAAATAAGGC | iPCR |
| iPCR_purL_1_25_B_MM14_F | TAAGCCTTACGGCTCAGTGCGTTTTAGAGCTAGAAATAGCAAGTTAAAATAAGGC | iPCR |
| iPCR_purL_3_201_C_F | GGAGTTTGCCTTGCGGGGCGGTTTTAGAGCTAGAAATAGCAAGTTAAAATAAGGC | iPCR |
| iPCR_purL_3_201_W_MM1_F | CGAGTTTGCCTTGCGGGGCGGTTTTAGAGCTAGAAATAGCAAGTTAAAATAAGGC | iPCR |
| iPCR_purL_3_201_B_MM5_F | CCTCATTGCCTTGCGGGGCGGTTTTAGAGCTAGAAATAGCAAGTTAAAATAAGGC | iPCR |
| iPCR_purL_3_201_B_MM9_F | CCTCAAACGCTTGCGGGGCGGTTTTAGAGCTAGAAATAGCAAGTTAAAATAAGGC | iPCR |
| iPCR_purL_3_201_B_MM14_F | CCTCAAACGGAACGGGGGCGGTTTTAGAGCTAGAAATAGCAAGTTAAAATAAGGC | iPCR |
| qPCR_dapA_F | GCGGTCATGGGGTTATTTCC | qPCR |
| qPCR_dapA_R | TGTGTAATGGCATCAGACGC | qPCR |
| qPCR_dapB_F3 | AGGTACGCTGAACCATCTCG | qPCR |

| | | |
|---|---|---|
| qPCR_dapB_R3 | ACGTTAACGCCAACGCTAAA | qPCR |
| qPCR_gdhA_F2 | CATTTGGCGGCAGTCTTATT | qPCR |
| qPCR_gdhA_R2 | TCCATCGCTTTTTCGATAGC | qPCR |
| qPCR_gltB_F2 | AAAACTACGCTGTCGGGATG | qPCR |
| qPCR_gltB_R2 | AGAGAGGAGAGGGCGATTTC | qPCR |
| qPCR_folA_F2 | GCCATACCTGGGAATCAATC | qPCR |
| qPCR_folA_R2 | TCCACTTCTGCGTCGATATG | qPCR |
| qPCR_thyA_F2 | TGAACTGCTGTGGTTTCTGC | qPCR |
| qPCR_thyA_R2 | GGTCGTTTTTCAGCTGGTTC | qPCR |
| qPCR_glyA_F4 | ATCGATCGTGCGAAAGAACT | qPCR |
| qPCR_glyA_R4 | GTTAACCGGAGAACCGTGAG | qPCR |
| qPCR_purN_F1 | TGACGCCTGTAAAACCAACA | qPCR |
| qPCR_purN_R1 | ATAAAACCAGCCAGCACGAC | qPCR |
| qPCR_purL_F1 | ACTTGAACGCCTGCTGAAAT | qPCR |
| qPCR_purL_R1 | GGTAACCTGCTGCCATTGTT | qPCR |
| qPCR_hcaT_F2 | TGGCACTGCTGACACTTCTC | qPCR |
| qPCR_hcaT_R2 | AGTCGCACTTTGCCGTAATC | qPCR |
| qPCR_gDNA_F1 | TATTTTGCGTTCCCGTTTGT | qPCR |
| qPCR_gDNA_R1 | CCCTCTATTCACCCGAAACA | qPCR |
| BC_iPCR_F_BC1 | CTTTCAGTTGCCTCGCTGTACTAATAATGG | pCRISPR3 Barcode Swap |
| BC_iPCR_F_BC2 | ATCATGGTTGCCTCGCTGTACTAATAATGG | pCRISPR3 Barcode Swap |
| BC_iPCR_F_BC3 | GCATGGGTTGCCTCGCTGTACTAATAATGG | pCRISPR3 Barcode Swap |
| BC_iPCR_F_BC4 | GTATGAGTTGCCTCGCTGTACTAATAATGG | pCRISPR3 Barcode Swap |
| BC_iPCR_F_BC5 | AGTCTAGTTGCCTCGCTGTACTAATAATGG | pCRISPR3 Barcode Swap |
| BC_iPCR_F_BC6 | CCTAGTGTTGCCTCGCTGTACTAATAATGG | pCRISPR3 Barcode Swap |
| BC_iPCR_UnivR_v2 | ACGGATCCCCACCTTC | pCRISPR3 Barcode Swap |
